# Co-catabolism of arginine and succinate drives symbiotic nitrogen fixation

**DOI:** 10.1101/741314

**Authors:** Carlos Eduardo Flores-Tinoco, Matthias Christen, Beat Christen

## Abstract

Biological nitrogen fixation emerging from the symbiosis between bacteria and crop plants holds a significant promise to increase the sustainability of agriculture. One of the biggest hurdles for the engineering of nitrogen-fixing organisms is to identify the metabolic blueprint for symbiotic nitrogen fixation. Here, we report on the CATCH-N cycle, a novel metabolic network based on co-catabolism of plant-provided arginine and succinate to drive the energy-demanding process of symbiotic nitrogen fixation in endosymbiotic rhizobia. Using systems biology, isotope labeling studies and transposon sequencing in conjunction with biochemical characterization, we uncovered highly redundant network components of the CATCH-N cycle including transaminases that interlink the co-catabolism of arginine and succinate. The CATCH-N cycle shares aspects with plant mitochondrial arginine degradation pathway. However, it uses N2 as an additional sink for reductant and therefore delivers up to 25% higher yields of nitrogen than classical arginine catabolism — two alanines and three ammonium ions are secreted for each input of arginine and succinate. We argue that the CATCH-N cycle has evolved as part of a specific mechanism to sustain bacterial metabolism in the microoxic and acid environment of symbiosomes. In sum, our systems-level findings provide the theoretical framework and enzymatic blueprint for the rational design of plants and plant-associated organisms with new properties for improved nitrogen fixation.

**Significance Statement:** Symbiotic bacteria assimilate nitrogen from the air and fix it into a form that can be used by plants in a process known as biological nitrogen fixation. In agricultural systems, this process is restricted mainly to legumes, yet there is considerable interest in exploring whether similar symbioses can be developed in non-legumes including cereals and other important crop plants. Here we present systems-level findings on the minimal metabolic function set for biological nitrogen fixation that provides the theoretical framework for rational engineering of novel organisms with improved nitrogen-fixing capabilities.

---

Nitrogen is a fundamental element of all living organisms and the primary nutrient that impacts crop yield (1, 2). Despite being highly abundant in the atmosphere, plants can only utilize nitrogen in reduced forms such as ammonium. More than 125 megatons of nitrogen are fixed annually by the industrial Haber-Bosch process into ammonium and applied to increase agricultural crop production (2). On the global scale, anthropogenic nitrogen delivered to the environment surpasses annual supplies by natural biological nitrogen fixation on land (3) leading to serious environmental impacts from climate change to the disruption of eco-systems and pollution of coastal waters.

Improving the ability of plants and plant-associated organisms to fix atmospheric nitrogen has inspired biotechnology for decades (4–6), not only for the apparent economic and ecological benefit that comes with the replacement of chemical fertilizers but also more recently for opportunities towards more sustainable agriculture and the potential to reduce greenhouse gas emissions. Attempts to transfer and improve nitrogenase genes clusters have to date focused largely on organisms such as *Escherichia coli* (7, 8). More recently the emerging field of synthetic biology provides an alternative approach to engineer designer nitrogenase gene clusters in bacteria (9–12). Despite these promising results, engineered organisms based on het erologous expression of nitrogenase genes have not yet come close to the efficiency of natural rhizobia-legume symbiosis systems (4). While the molecular mechanism of the nitrogenase reaction has been resolved with atomistic detail (13–16), the precise mechanism how the metabolism between plants and bacteria becomes entangled to sustain the energy-intensive process of nitrogen fixation has remained an open question.

The current model of nutrient exchange in rhizobia-legume symbiosis postulates that, in exchange for fixed nitrogen, the plant provides dicarboxylic acids such as succinate, which is metabolized through the tri-carboxylic acid (TCA) cycle to generate ATP and reduction equivalents needed for the nitrogenase reaction (17–19). However, multiple lines of evidence argue against a simple exchange of succinate for ammonium during symbiosis (20, 21). While the absence of oxygen promotes nitrogenase activity, it also inhibits TCA-mediated catabolism of succinate due to redox inhibition of key TCA-enzymes including citrate synthase, isocitrate dehydrogenase, and 2-oxoglutarate dehydrogenase (22, 23). Thus, the TCA cycle probably operates below its full aerobic potential. Furthermore, if the metabolism of symbiotic nitrogen-fixing bacteria is based exclusively on the provision of succinate, then the bacterial nitrogen requirement must be covered solely by nitrogenase. However, nitrogen-fixing root-nodule bacteria (termed bacteroids), do not self-assimilate but rather secrete large quantities of ammonium (24–26) suggesting that the plant provides the bacteroids with a nitrogen-containing nutrient to cover their nitrogen needs. Finally, the degradation product of succinate in the TCA cycle is carbon dioxide. However, it has been reported that nitrogen-fixing bacteroids also secrete the amino acids alanine and aspartate (27–29). Here, we report on the CATCH-N cycle based on the co-catabolism of plant-provided arginine and succinate as part of a specific metabolic network to sustain symbiotic nitrogen fixation. Using isotope-labeled mass-spectrometry analysis in *Bradyrhizobium diazoefficiens* in conjunction with *in planta* transposon-sequencing analyses and enzymatic reaction network characterization in *Sinorhizobium meliloti*, we uncovered the principle of metabolic entanglement leading to the nitrogen-fixing symbiosis between plants and bacteria. Collectively, we demonstrate that the CATCH-N metabolism is governed by highly redundant functions comprised of at least 10 transporter systems and 23 enzymatic functions. In sum, our systems-level findings provide the theoretical framework and enzymatic blueprint for the optimization and redesign of improved symbiotic nitrogen-fixing organisms.

## Results

### Evidence for plant-provided arginine driving nitrogen fixation in rhizobia-legume symbiosis

We asked what the identity of the postulated nitrogen-containing nutrient could be. Since the plant must provide the compound in sufficient quantities, we reasoned that an amino acid might be a likely candidate. Based on the finding that nitrogen-fixing bacteroids utilize succinate and secrete the amino acids alanine and aspartate (27–30), we concluded that the plant-provided compound must comprise at least two nitrogen atoms to enable two consecutive transamination reactions. The first nitrogen is used for transamination of the ketoacid derived from succinate while the second nitrogen atom is utilized for transamination of the ketoacid derived from the plant-provided compound.

Six out of the twenty natural amino acids (Arg, His, Lys, Gln, Asn, and Trp) contain two or more nitrogen atoms and thus are likely candidates. Thereof, His, Lys and Gln can be excluded because their degradation involves a compulsory 2-oxoglutarate dehydrogenase step, which is subjected to redox inhibition and disfavoured under microoxic conditions (31, 32). Furthermore, we also excluded Trp and Asn because their catabolism enters the TCA at the level of pyruvate and oxaloacetate respectively, which limits energy metabolism within a partially operating TCA cycle. Based on these theoretical considerations, we postulated that the remaining amino acid arginine is a likely candidate for the nitrogen-containing compound provided upon symbiosis.

### The co-feeding of arginine and succinate stimulates nitrogenase activity

To probe whether arginine functions as cosubstrate to drive symbiotic nitrogen fixation, we assayed nitrogenase activity of mature bacteroids from *B. diazoefficiens* (strain 110 spc4) and *S. meliloti* (strain CL 150) in presence of nodule crude extracts and upon supplementation of succinate and arginine (materials and methods). The addition of nodule crude extracts to isolated bacteroids resulted in strong stimulation of nitrogenase activity, supporting the idea that plant provided nutrients are necessary for symbiotic nitrogen fixation. While the stimulation of nitrogenase only poorly occurred in the presence of succinate as the sole nutrient, we found that the addition of arginine stimulated nitrogenase activity in *B. diazoefficiens* and *S. meliloti* to 50% ± 4% and 116% ± 2% respectively as compared to nodule extracts.Furthermore, the co-feeding of arginine in combination with succinate restored nitrogenase activity to the same extent as nodule extracts (92% ± 6% and 92% ± 6%) for *B. diazoefficiens* (Fig. 1A) and *S. meliloti* bacteroids respectively. Therefore, we concluded that the co-feeding of arginine and succinate is sufficient to stimulate nitrogenase activity in bacteroids.

**Fig. 1.**
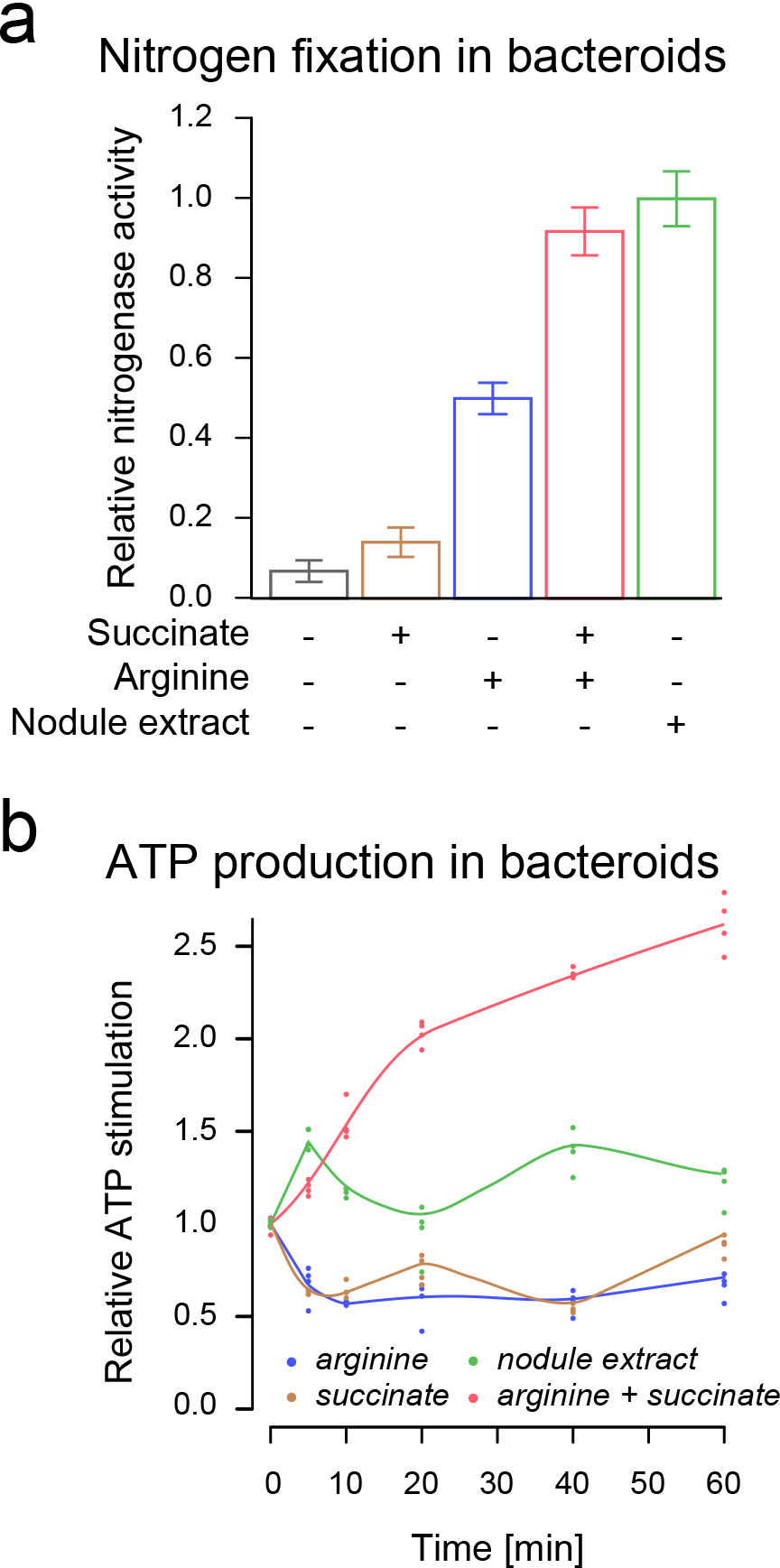
The co-feeding of arginine and succinate promotes nitrogen fixation and ATP production in isolated bacteroids. (**a**) Substrate-dependent nitrogenase activity in isolated *B. diazoefficiens* bacteroids upon supplementation of succinate, arginine, and nodule extract. Data represents the mean and standard error of the mean. (**b**) ATP stimulation in isolated *B. diazoefficiens* bacteroids upon supplementation of succinate, arginine, and nodule extract.

The nitrogenase enzyme complex catalyzes one of the most energy-consuming enzymatic reactions found in nature with 16 ATP molecules and 8 low-potential electrons required for the reduction of a single nitrogen molecule. Nitrogenase is irreversibly inactivated in the presence of oxygen, which restricts the reduction of atmospheric nitrogen to low-oxygen conditions. Thus, to support nitrogen fixation, bacteroids must produce substantial amounts of ATP under microoxic conditions. The finding that succinate as the sole nutrient did not result in nitrogenase stimulation suggested that succinate catabolism via the TCA cycle does likely not generate sufficient ATP to support efficient nitrogenase reaction.

To measure the ATP level produced in isolated *B. diazoefficiens* bacteroids, we quantified the increase in intracellular ATP through ATP-dependent luciferase assays (materials and methods). In agreement with the absence of an operational TCA cycle, we observed that the addition of succinate alone failed to stimulate ATP production. In contrast, we found that co-feeding of succinate together with arginine caused an increase from 1.53 ± 0.04 to 4.03 ± 0.09 attomole ATP per cell corresponding to a 2.61 ± 0.03 fold stimulation (Fig. 1B). In sum, these findings suggest that co-catabolism of arginine and succinate supports biological nitrogen fixation in *B. diazoefficiens* and *S. meliloti*.

### Isotope tracing experiments reveal the presence of three parallel arginine degradation pathways

To gain further insights into the structure and dynamics of the arginine degradation network operating during nitrogen fixation, we performed stable isotope labelling studies with *B. diazoefficiens*. We incubated isolated bacteroids under stringent microoxic conditions with ^13^C arginine in the presence of unlabelled succinate and quantified isotope labelling pattern of arginine degradation intermediates by LC-MS/MS (Table 1). Upon the addition of ^13^C arginine, we observed a rapid increase in the labeled intracellular arginine pool (99.43% ^13^C), demonstrating active arginine transport into nitrogen-fixing bacteroids.

**Table 1.** Arginine catabolism in *B. diazoefficiens* bacteroids fed with ^13^C arginine and unlabeled succinate.

| Metabolite | Fractional labeling (%)<sup>a</sup> |
| --- | --- |
| Arginine (ARG) | 99.43 ± 0.10 |
| Citrulline (CIT) | 60.22 ± 5.26 |
| Ornithine (ORN) | 81.25 ± 5.01 |
| Proline (PRO) | 23.19 ± 3.58 |
| Glutamate (GLU) | 7.67 ± 0.82 |
| 4-guanidinobutanoate (GBA) | 90.40 ± 3.03 |
| 4-aminobutanoate (GABA) | 6.26 ± 1.06 |

<sup>a</sup> $^{13}\text{C}$ Fractional labeling after 150 min incubation with $^{13}\text{C}$ L-arginine. Shown is the average and the standard error of the mean (SEM).

Upon further incubation, we found ^13^C isotope labels in key intermediates of multiple arginine degradation pathways steadily increasing. After 150 minutes, ornithine, proline, and glutamate, the intermediates of the classical arginase-mediated degradation pathway, were labeled with 81.25%, 23.19% and 7.67% ^13^C (Table 1). Also, citrulline, which represents the first step of the arginine deiminase (ADI) pathway, was labeled with 60.22% ^13^C. This finding suggests that bacteroids employ the ADI pathway for anaerobic, enzyme-coupled ATP production by the enzyme carbamate kinase.

Furthermore, we also observed fractional labelling of 90.40% for 4-guanidinobutanoate (GBA) and 6.26% for 4-aminobutanoate (GABA) further suggesting the presence of a functional arginine transaminase (ATA) pathway operating in bacteroids (Table 1) that yields alanine. The observation that bacteroids posses an ATA pathway was intriguing because it provided a functional link between arginine degradation and alanine secretion, which was previously reported as part of the metabolite exchange occurring during symbiotic nitrogen fixation (30). These findings suggest that at least three independent arginine degradation pathways operate simultaneously in nitrogen-fixing *B. diazoefficiens* bacteroids. Collectively, arginine degradation results in the production of alanine and ammonium independent from the nitrogenase reaction.

### Transposon sequencing reveals symbiosis genes involved in the uptake and catabolism of arginine

To gain further insights into the gene sets and enzymatic functions responsible for uptake and degradation of arginine, we conducted a functional genetic screen using transposon sequencing (TnSeq) (33, 34). TnSeq measures genome-wide changes in transposon insertion abundance prior and after subjecting large mutant populations to selection regimes (35) and allows systems-level definition of conditional essential gene sets for a given environment (36, 37). We reasoned that TnSeq provides a unique opportunity to identify specific arginine transport and degradation genes that become essential upon engagement in symbiosis.

We choose *S. meliloti-Medicago truncatula* as the rhizobia-legume symbiosis system, because supernodulating *M. truncatula lss* plants (38) provided a high frequency of nodules increasing the resolution of the TnSeq analysis. In total, we infected 4,500 *M. truncatula lss* plants with a high-density *S. meliloti* transposon mutant library of 750,128 unique Tn*5* insertions (Fig. 2A). Six weeks post-inoculation, we recovered 99,623 unique Tn*5* mutants from 375,000 root nodules. By comparing the TnSeq dataset obtained from plant infection assays and input transposon mutant libraries, we mapped a set of 977 symbiosis genes corresponding to 15.71% of the tripartite 6.7-megabase (Mb) genome (Data SI, Materials and methods). Thereof, 435 genes were located on the chromosome, 295 on pSymA and 247 on pSymB indicating that all three replicons of *S. meliloti* contribute to symbiotic nitrogen fixation (Fig. 2B).

**Fig. 2.**
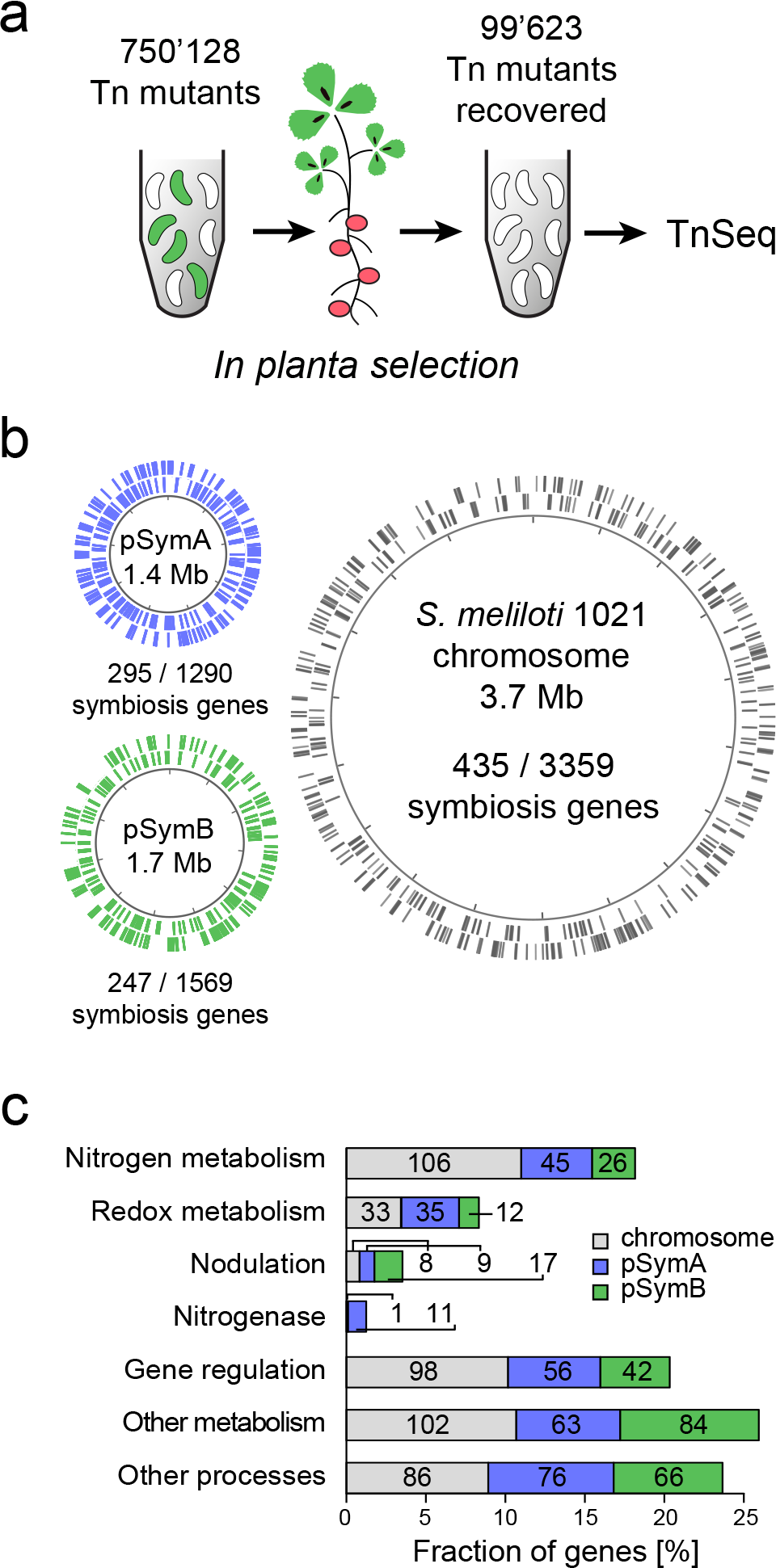
The symbiosis genome of *S. meliloti* revealed by transposon sequencing (TnSeq). (**a**) Schematic representation of the plant infection screen that was used to map the *S. meliloti* symbiosis genome. Tn*5* transposon mutant pools were selected for their ability to establish symbiosis with *M. truncatula*. After selection, Tn*5* mutants recovered from root nodules were identified by TnSeq. (**b**) Genome map visualizing the distribution of essential symbiosis genes among the three *S. meliloti* replicons. Symbiosis genes are plotted as lines on the chromosome (grey) and the megaplasmids pSymA (blue) and pSymB (green). (**c**) Functional classification of essential symbiosis genes located on the chromosome (grey), pSymA (blue) and pSymB (green)

Functional classification revealed that the large majority of symbiosis genes comprise cellular functions such as metabolism (507 genes, 51.89%), gene regulation (196 genes, 20.06%) and other cellular processes (228 genes, 23.34%) (Fig. 2C, Data SI). The identified gene set included well-characterized symbiosis factors involved in nodulation (34 genes, 3.48%) as well as functions associated with the nitrogenase enzyme complex (12 genes, 1.23%). Collectively, a set of 177 symbiosis genes, corresponding to over one-third of the 507 metabolic symbiosis genes (34.91%), was associated with nitrogen metabolism including genes for the transport (59 genes), biosynthesis (78 genes) and degradation (40 genes) of amino acids and other nitrogen-containing compounds. While only 3 out of 78 essential biosynthesis genes (3.85%) were involved in the synthesis of arginine, we found a large fraction of 18 out of 59 essential transport genes (30.51%) and 22 out of 40 essential catabolic genes (55.00%) annotated as being involved in the uptake and catabolism of arginine and its derivatives. In sum, these findings highlight that the provision of arginine and its consecutive degradation is of fundamental importance to drive symbiotic nitrogen fixation in legumes.

### Tnseq identifies multiple arginine transport systems mediating acid tolerance

Among the 18 transport genes essential for symbiosis, we found two arginine and four putrescine ABC transport systems that we named *artABCDE* (SMc03124-28) for arginine transporter, *satABC* (SMa2195-97) for symbiotic arginine transporter and *potFGHI* (SMc00770-3), *potABCD2* (SMa0799-803), *potABCD3* (SMa0951-3) and *potABCD4* (SMa2203-9) for the putrescine uptake systems (Data SI). In addition, two arginine/agmatine antiporter genes *adiC* (SMa0684) and *adiC2* (SMa1668) encoded on pSymA were also essential during symbiosis. Interestingly, all identified transport systems participate in urease, and ADI pathways that mediate acid tolerance. Cross-comparing expression profiles using previously published RNA-seq data sets (39), we found that *artABCDE* was the only transport system to be constitutively expressed during all stages of symbiosis, while the expression of all other transporters was specifically induced during development into nitrogen-fixing compartments (symbiosomes).

To gain further insights, we searched for additional symbiosis genes related to acid tolerance and indeed found multiple essential components in the TnSeq dataset (Data SI). From the urease pathway, we identified two arginases (*argI1* and *argI2*) and the urease components *ureA* and *ureE*. From the ADI system, we found the *arcABC* operon to be essential for symbiosis. Both systems catalyze the conversion of arginine into ornithine leading to the production of ammonia as part of the acid tolerance mechanism. Furthermore, the ADI system also provides ATP via the enzymatic step of ornithine carbamoyltransferase *arcB* (40). Interestingly, two additional copies of ornithine carbamoyltransferase were also essential (*argF1*, and *arcB2*), emphasizing the importance of genetic redundancy in ADI dependent ATP synthesis during symbiosis. The urease and ADI acid tolerance mechanisms rely on the efflux of ammonium (41). Indeed, the ammonium efflux pump encoded by *amtB* was among the top-ranked symbiosis genes. These findings underscore the importance of ammonium secretion as a compulsive property of bacteroids independent of the nitrogenase reaction.

### Arginase gene deletions show nitrogen starvation phenotypes during plant infection assays

To validate the importance of the identified arginine-dependent acid tolerance systems for symbiosis, we constructed a panel of deletion mutants of the urease and ADI pathways and assessed nitrogen starvation phenotypes during plant infection assays. Out of the 8 mutants evaluated, all displayed symbiotic defects. On the level of the arginine transport systems, we found that *artABCDE* and *satABC* showed a reduction in nitrogenase activity of 47.06% ± 7.27% and 55.45% ± 10.99%. Similarly, gene deletions in the urease pathway such as the arginases mutants *argI1* and *argI2* exhibited a reduction of 71.18% ± 5.21% and 70.97 ± 5.16% for single deletions and 80.89% ± 3.15% for the double deletion mutant. Deletion of the urease *ureGFE* and the ammonium efflux system *amtB* resulted in a 64.68% ± 5.50% and 80.90% ± 4.81% reduction in nitrogenase activity.

Plants inoculated with the *argI1,argI2* single and double deletion mutant harbored a typical phenotype of nitrogen starvation. The aerial part of infected plants was smaller than those inoculated with WT strain (Fig. 3A, S1). Nodules induced by the *argI1,argI2* double deletion mutant displayed the yellowish color of non-functional *M. truncatula* nodules (Fig. 3B). Furthermore, observations of *argI1,argI2* nodule sections by scanning electron microscopy showed that nodules were hollow and remaining bacteroids exhibit aberrant cell morphology (Fig. 3C, S2). Collectively, our results suggest that the identified arginine-dependent acid tolerance system is a prerequisite for the faithful establishment of symbiosis. (SMa2203-9) for the putrescine uptake systems (Data SI).

**Fig. 3.**
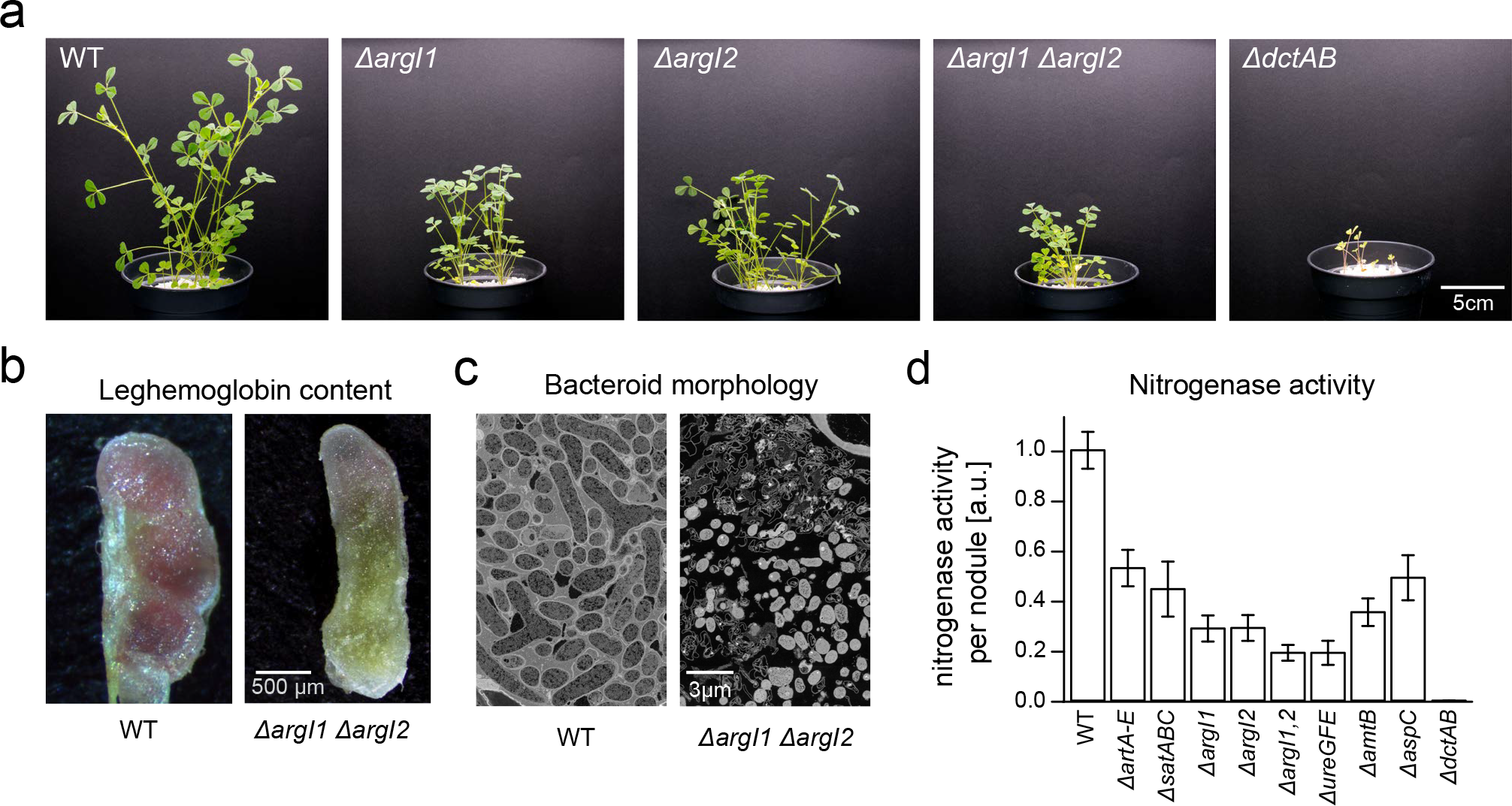
Assessment of nitrogen starvation phenotypes of *M. truncatula* upon infection with *S. meliloti* mutants impaired in arginine transport and catabolism. (**a**) The aerial part of *M. truncatula* upon infection with *S. meliloti* Δ*argI*1, Δ*argI*2, Δ*argI*1,2 and the dicarboxylate transport mutant Δ*dctAB* were smaller than those inoculated with the wild type strain, highlighting the importance of arginine catabolism for nitrogen fixation (**b**) Cross sections of nodules bearing *S. meliloti* WT or arginine catabolism mutant Δ*argI1* Δ*argI2* reveals the yellowish color of non-functional nodules induced by the Δ*argI1* Δ*argI2* double deletion mutant. (**c**) Bacteroid ultrastructure across nodule sections determined by scanning electron microscopy indicates the presence of hollow nodules with aberrant cell morphology in the Δ*argI1* Δ*argI2* double deletion mutant defective in arginine catabolism.(**d**) Nitrogenase activity in *M. truncatula* nodules inoculated with *S. meliloti* strains defective in arginine catabolism. Data points are the mean of at least 30 plants measured after 8 weeks post inoculation; error bars indicate standard error of the mean

### Identification of AspC as an arginine:pyruvate transaminase

In our isotope labelling studies with bacteroids, we detected fractional labelling of 90.40% for GBA suggesting the presence of a functional ATA pathway. However, in the *S. meliloti* genome, corresponding genes have not been assigned. Among the essential symbiosis genes identified by our TnSeq analysis, we found *aspC* (SMc02262) annotated as an aspartate transaminase, which is specifically expressed in nitrogen-fixing bacteroids (39). We constructed a clean deletion mutant of *aspC* and found that the absence of AspC indeed causes a nitrogen starvation phenotype during plant infection assays. The aerial part of plants infected with the *aspC* deletion mutant were smaller than those inoculated with wildtype strain and nitrogenase activity was reduced by 50.91 % ± 9.00% (Fig. 3D, Table S1) highlighting the importance of *aspC* for nitrogen fixation.

AspC shares 40% sequence homology to AruH, the arginine:pyruvate transaminase from *Pseudomonas aeruginosa* (42, 43). When we characterized the enzymatic properties of AspC by mass spectrometry, we found transaminase activity to pyruvate from arginine, agmatine, ornithine, and putrescine (Table S2) to yield alanine. Enzyme studies with *ilvB1*, the gene encoded downstream of *aspC*, further provided evidence that the *S. meliloti* ATA pathway differs from *P. aeruginosa*. Sequence homology with AruI from *P. aeruginosa* suggested that IlvB1 is a 5-guanidino-2-oxo-pentanoate (GOP) decarboxylase. However, assays with purified IlvB1 revealed that the *S. meliloti* enzyme has 7-fold higher decarboxylase activity with 5-amino-oxopentanoate (AOP). We concluded that both AspC and IlvB1 are promiscuous enzymes accepting multiple substrates. Although *aspC* was among the symbiosis genes, the deletion mutant retained partial nitrogenase activity. Also, *ilvB1* was not present within the group of symbiosis genes. This suggests that genetic redundancies among transaminase and decarboxylase genes exist and partially mask mutant phenotypes complicating the genetic identification of additional network components.

### At least 16 redundant enzymes participate in the arginine transamination network

To further dissect possible redundant components, we performed a functional homology search for enzyme candidates known to be expressed during the nitrogen-fixing bacteroid stage. Conversion from arginine into succinate proceeds by a series of six consecutive enzymatic reaction steps. The four steps upstream of GABA comprise transamination, decarboxylation, ureohydrolase and dehydrogenase reactions. In the lower part of the reaction network, a linear pathway through transamination and subsequent dehydrogenase steps leads from GABA to succinate (Fig. 4A). Upon heterologous expression and protein purification, we biochemically profiled a panel of 16 candidate enzymes. In addition to the arginine deiminases ArcA1 and ArcA2 and the arginase ArgI1, we found two agmatinase (ArgI2 and SpeB) and one ureohydrolase (SpeB2) acting on 4-guanidinobutyraldehyde (GBL), GOP and GBA (Fig. 4B). The highest level of pathway redundancy resides on the level of dehydrogenases. Besides the five known GabD1-5 proteins from *S. meliloti*, we identified four additional isoenzymes Gab6-9. Thereof GabD6 and GabD7 share a dehydrogenase profile identical to GabD1 for 4-aminobutyraldehyde (ABL), succinic semialdehyde (SSA) and GBL. GabD8 and GabD9 exhibited substrate specificity for GBL (Fig. 4B). Furthermore, on the level of pyruvate transaminases, we identified three additional enzymes (DatA, AatB, and ArgD) that exhibited substrate preferences for or nithine, putrescine and agmatine and two enzymes (GabT2, and GabT3) with preferences for GABA (Fig. 4B). Similarly, we also profiled two additional decarboxylase enzymes (OdcA, and OdcB) that either catalyzed the decarboxylation of ornithine or AOP (Fig. 4B). Collectively, these results demonstrate the presence of a highly redundant network mediating arginine catabolism in *S. meliloti*.

**Fig. 4.**
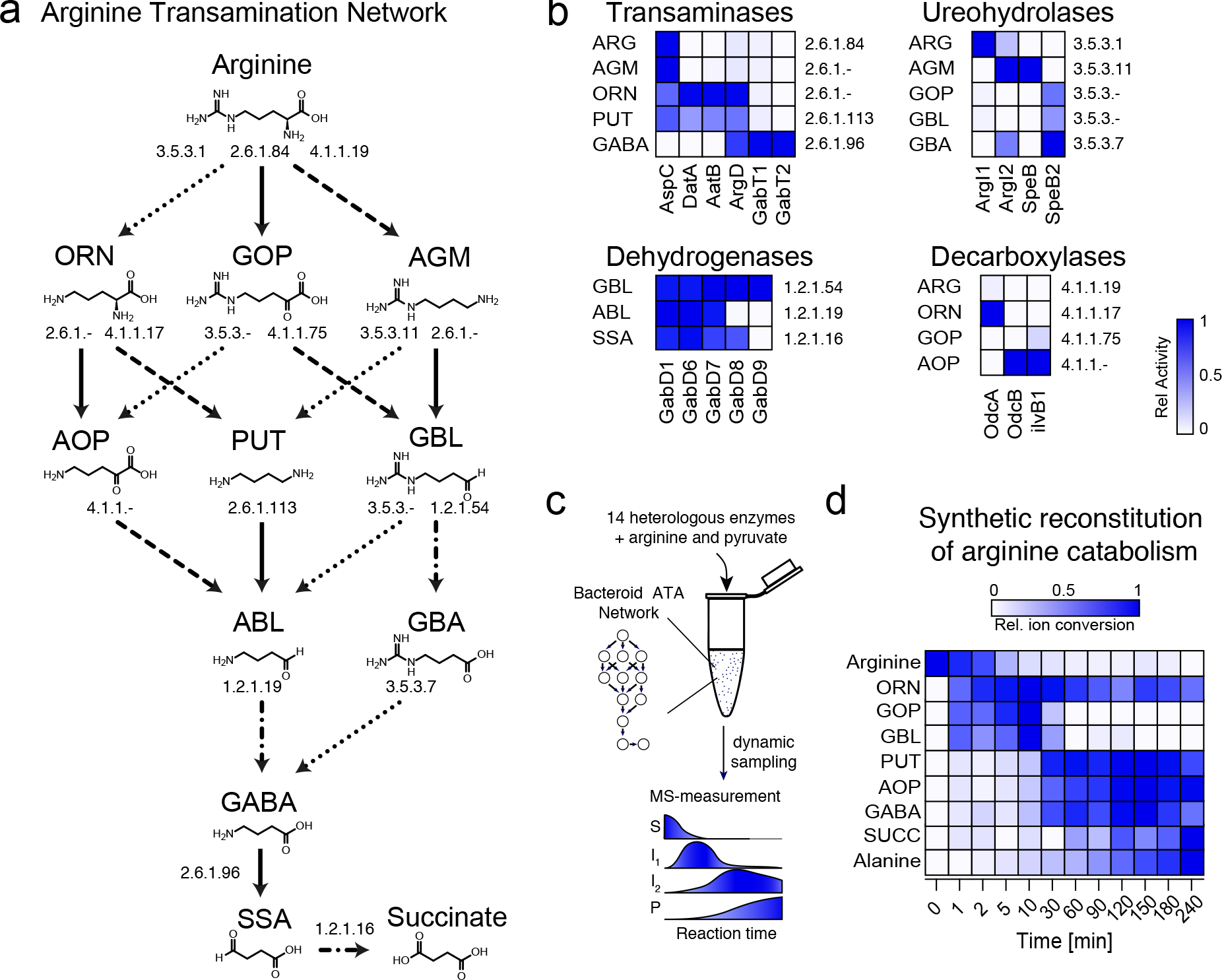
Reaction scheme, enzyme profiling, and reconstitution of the catabolic arginine transamination network operating in nitrogen-fixing bacteroids. (**a**) Schematic representation of the arginine-transamination reaction network than leads from arginine to succinate. Enzymatic reaction steps are indicated by arrows and annotated with corresponding enzyme commission (EC) numbers. Leftward facing dotted arrows represent ureohydrolase, rightward facing dashed arrows decarboxylases, downward-pointing solid arrows aminotransferases, and dashed and dotted arrows dehydrogenase enzyme reaction steps. The names of the metabolites are indicated above their chemical structures according to the following abbreviations: arginine (ARG), ornithine (ORN), 5-guanidino-2-oxo-pentanoate (GOP), agmatine (AGM), 5-amino-oxopentanoate (AOP), putrescine (PUT), 4-guanidino-butyraldehyde (GBL), 4-aminobutyraldehyde (ABL), 4-guanidinobutanoate (GBA), 4-aminobutanoate (GABA), succinic semialdehyde (SSA), and succinate (SUCC). (**b**) Heat map representing enzymatic activities participating in the arginine transamination network with relative enzyme activities highlighted as a color map. Rows are annotated on the left with the abbreviated names of the substrates and on the right with the corresponding EC numbers. Columns are annotated by corresponding enzyme names. (**c**) Recombinant enzymes involved in arginine catabolism were combined into a single reaction mixture using arginine and pyruvate as substrates. Substrate (S) consumption into the products (P) succinate and alanine, including their intermediates (I_1_,I_2_), was determined along the time series using LC-MS/MS. (**d**) The relative metabolite concentration along the reaction time-course is highlighted as a color map.

### Synthetic reconstitution of the arginine transamination network that operates in nitrogen-fixing bacteroids

We reconstituted the reaction system *in vitro* from a set of 14 enzymes and followed the conversion of arginine and pyruvate by mass spectrometry (Fig. 4C). We observed that 90% of the arginine was rapidly metabolized within 30 minutes. As expected, ornithine and GOP appeared as the first intermediates and then concomitantly decreased with the appearance of the second level of intermediates putrescine, AOP and ABL that ultimately converted into GABA, SSA, and succinate. During the process, alanine steadily increased demonstrating that pyruvate transamination couples the conversion of arginine into succinate (Fig. 4D). In sum, these findings demonstrate the synthetic reconstruction of the transamination network that permits the co-catabolism of arginine and succinate.

### The catabolism of succinate and arginine is interlinked

Since the ATA network consumes two equivalents of pyruvate but generates only a single equivalent during its operation, we reasoned that the network strictly depends on the provision of additional pyruvate, which must be formed by simultaneous co-catabolism of succinate. We concluded that the degradation of arginine and succinate are mutually coupled and can only take place if plants provide both nutrients in equal stoichiometries. Indeed, single deletion in the succinate transporter DctAB abrogates succinate uptake and thereby also prevents the co-catabolism of arginine, resulting in a fix minus phenotype. On the other hand, the uptake of arginine is controlled by multiple redundant transporters. Nevertheless, single deletions in *artABCDE* and *satAB* arginine transporter show a partial symbiosis phenotype. Thus, transamination enforces a strict co-catabolism of succinate and arginine but also provides an elegant solution to maintain a partial TCA cycle under stringent microoxic and acidic conditions. We termed the entangled catabolic network CATCH-N cycle for C4-dicarboxylate Arginine-Transamination Co-catabolism under acidic (H^+^) conditions to fix Nitrogen (Fig 5).

**Fig. 5.**
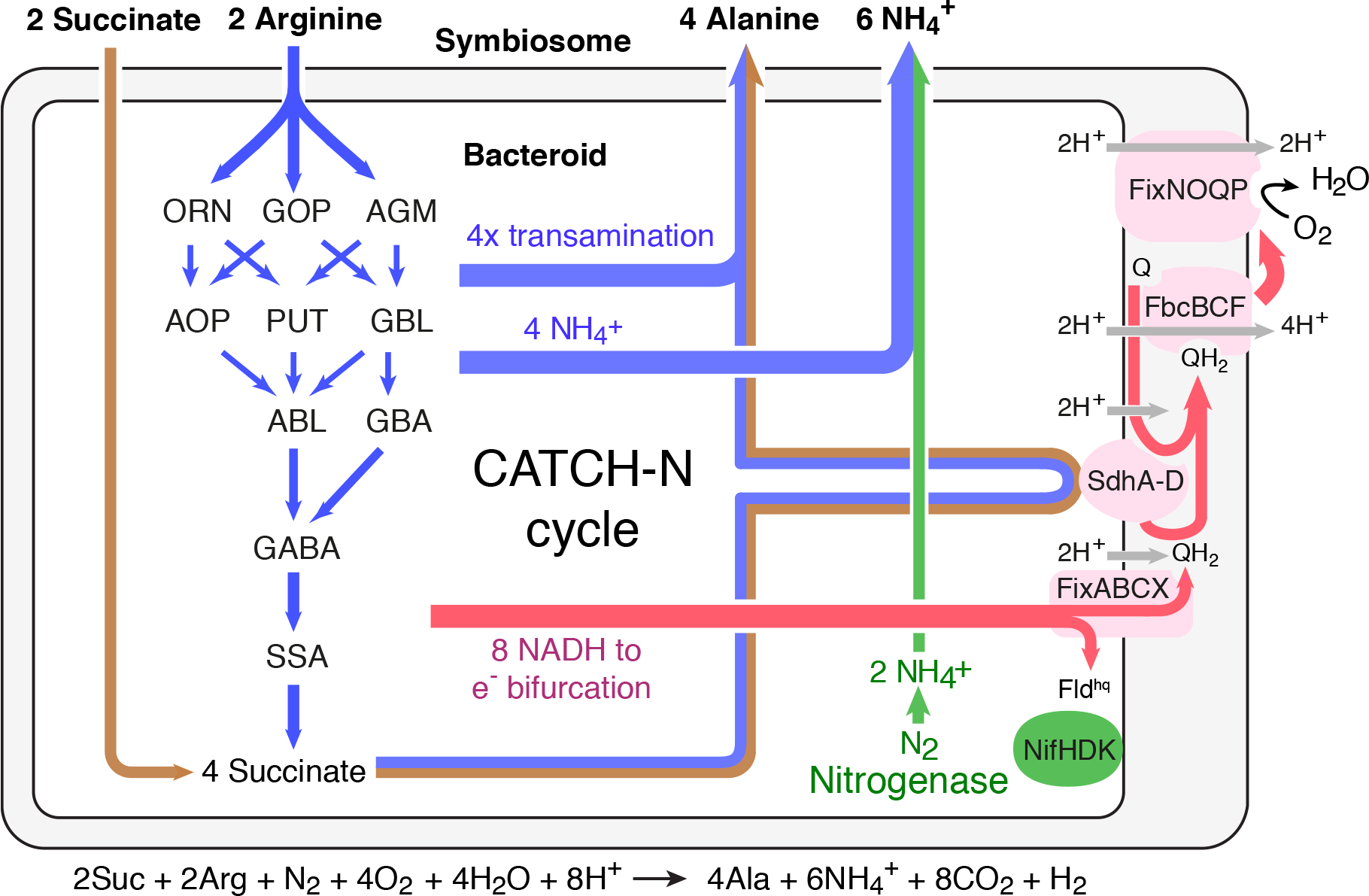
Model of the CATCH-N cycle operating in N_2_-fixing bacteroids. Arginine (blue) and succinate (brown) are co-feed to bacteroids in equimolar ratio. Co-catabolism is interlinked through an arginine-pyruvate transamination step yielding alanine. Alternatively, an arginine-oxaloacetate transamination step yields aspartate. Enzymatic conversion of arginine releases ammonium (blue) independent from the nitrogenase reaction (green). NADH (magenta) produced through the co-catabolism is regenerated over a bifurcated electron transport chain onto terminal acceptors oxygen and nitrogen

### A bifurcated electron transport chain operates during nitrogen fixation

We reasoned that the operation of the CATCH-N cycle provides significant amounts of NADH as well as QH_2_. Under aerobic conditions, NADH is regenerated by the electron transport chain, which includes proton-pumping enzymes known as complex I, III and IV. However, upon symbiosome acidification, the driving force of complex I is likely no longer sufficient to sustain proton translocation against the increased pH gradient impairing the conversion of NADH into QH_2_. Also, oxygen partial pressure within symbiosomes is too low to operate the aerobic version of complex IV and bacteroids induce expression of a high-affinity cytochrome cbb3 oxidase complex FixNOQP1-3 (44–46). Recently, the electron bifurcating FixABCX protein complex has been proposed to serve as the alternative entry point for low-potential (‘high-energy’) electrons from NADH (47, 48) thereby bypassing the impaired complex I.

Based on these findings, we devised a model that restores electron flow from NADH to QH_2_ by electron bifurcation to nitrogenase and the high-affinity terminal oxidase (Fig 5). If this is the main pathway that permits regeneration of NADH, then all components must be essential in symbiosis. Indeed, we found genes encoding for components of the nitrogenase *nifHDK*, the electron bifurcation complex *fixABCX* and the alternative complex IV *fixNOQP*1-3 among the top-ranked symbiosis genes in the TnSeq dataset (Supplementary Dataset SI). These genetic evidences suggested that a bifurcated electron transport chain operates during nitrogen-fixing symbiosis.

### Estimation of the ATP balance of the bifurcated electron transport chain

The endergonic branch of the electron bifurcation reaction generates low-potential reducing equivalents in the form of flavodoxin hydroquinone (Fld^hq^) for nitrogenase catalysis (Fig 5). For every Fld^hq^ the nitrogenase consumes two additional ATP molecules. However, the exergonic branch of the electron bifurcation reaction translocates only three protons corresponding to a single ATP that is generated per electron passing from QH_2_ to coenzyme Q onto oxygen. Thus, the electron bifurcation of each NADH appears to be associated with a net loss of one ATP. In addition to NADH, the CATCH-N cycle also provides QH_2_ via succinate dehydrogenase (Fig 5). Thereby, up to two ATP are generated per QH_2_ passing its electrons onto oxygen. Accordingly, a bifurcated electron transport chain in combination with an active succinate dehydrogenase complex delivers a net gain in ATP, provided that catabolism generates NADH to QH_2_ in a 2:1 ratio. Contrary to this criterion, the complete TCA operating with malate or succinate provides a higher ratio of NADH to QH_2_ of 5:1 and 5:2 and, thus, inevitably results in a net loss of ATP.

In addition to proton-motive force-dependent ATP synthesis, several metabolic cycles including the CATCH-N and TCA provide additional ATP through enzyme-coupled synthesis. Furthermore, proton gradients are not exclusively generated by the proton expelling complexes of the electron transport chain, but moreover can also be established through proton consuming enzymatic reactions in the cytosol, including the production of ammonia and decarboxylation reactions implemented in the CATCH-N cycle. Based on these considerations, we calculated the net proton consumption for several theoretical nitrogen fixation cycles (CATCH_N1-4) that start with arginine and succinate and feature dedicated transamination reactions that channel the TCA products pyruvate and oxaloacetate into the corresponding amino acids alanine and aspartate (Supplementary Information). In contrast to earlier approaches (49), we did not restrict our design to previously annotated pathways but rather included all enzymatic activities of the arginine transamination network identified during our biochemical enzyme studies. To evaluate the feasibility of these theoretical cycles, we estimated the net gain in ATPper N2 molecule converted. Our calculations show that in terms of ATP production, the CATCH-N cycles are generally more energy-efficient than a stand-alone TCA cycle operating on succinate or malate (Table 2) and were also more energy efficient than the core sequence of other naturally existing arginine degradation pathways, such as the arginine-pyruvate-glutamate super-pathway (Supplementary Information). As an example, two of our CATCH-N cycles generated on average over 18 fold more ATP per N2 converted into ammonia as compared to the core sequence of the TCA cycle operating with succinate as the sole substrate. In sum, these calculations establish a new conceptional framework to understand and engineer symbiotic nitrogen-fixing organisms with futureperspectives for agriculture.

**Table 2.** Comparison of selected metabolic pathways for N_2_ fixation.

| Pathway | Substrates | Products | Reaction steps | Sum NADH | Sum QH<sub>2</sub> | O<sub>2</sub> consumed per N<sub>2</sub> | Influx H<sup>+</sup> | ATP net gain |
| --- | --- | --- | --- | --- | --- | --- | --- | --- |
| CATCH-N1 | 2Suc, 2Arg | 2Ala | 9 | 8 | 4 | -4 | 56 (48+8) | 2.80 |
| CATCH-N2 | 2Suc, 2Arg | 2Asp | 9 | 8 | 4 | -4 | 52 (48+4) | 1.60 |
| CATCH-N3 | 2Mal, 2Arg | 2Ala | 9 | 8 | 1 | -3 | 44 (36+8) | -0.80 |
| CATCH-N4 | 2Mal, 2Arg | 2Asp | 9 | 8 | 1 | -3 | 40 (36+4) | -2.00 |
| TCA | 1.6Mal | CO<sub>2</sub> | 11 | 8 | 1.6 | -2.8 | 38.8 (33.6+5.2) | -2.76 |
| TCA | 1.6Suc | CO<sub>2</sub> | 11 | 8 | 3.2 | -3.6 | 48.8 (43.2+5.2) | 0.12 |

<sup>a</sup> Shown are the numbers of substrate, reaction steps and the numbers of NADH, QH<sub>2</sub> and oxygen molecules required (negative numbers) or generated (positive numbers) during conversion of N<sub>2</sub> into two molecules of ammonia. The influx in H<sup>+</sup> for each cycle is listed with O<sub>2</sub>-dependent proton translocation and O<sub>2</sub>-independent proton consumption by enzymatic reactions listed in brackets. An H<sup>+</sup>/ATP ratio of 3.33 was assumed for estimating the gain in ATP per N<sub>2</sub> converted.

## Discussion

Here, we report on the CATCH-N cycle operating on the co-feeding of arginine and succinate as part of a specific metabolic network that drives the process of symbiotic nitrogen fixation. The CATCH-N cycle shares aspects with the plant mitochondrial arginine degradation pathway (50, 51), however, it delivers up to 25% higher yield in nitrogen in the form of two alanines and three ammonium secreted for each co-feed arginine and succinate. Thus, from the plant’s perspective, the CATCH-N cycle multiplies the nitrogen releasing capacity of arginine. On the level of bacteroids, the CATCH-N cycle provides an elegant solution for maintaining an active respiratory chain under the highly acidic and microoxic conditions present within the lysosomal compartment of the symbiosome. Thus, the CATCH-N cycle also functions as an effective mechanism to promote the survival of bacteroids within infected plant cells. Equimolar arginine and succinate serve as substrates and a molar ratio of nitrogen to the oxygen of 1:4 is required to operate the CATCH-N cycle. Therefore, nitrogen-fixation still depends on oxygen as terminal acceptor, while harnessing elementary nitrogen as the second electron acceptor for reducing equivalents generated by the metabolism. Also, the CATCH-N cycle requires a constant flux of 8 protons into the symbiosome to maintain the pH balance of the reaction. These protons must be translocated by the action of plant ATPases as part of the symbiosome acidification process. Thus, the operation of the CATCH-N cycle depends on the presence of an active plant metabolism. From the nitrogen balance standpoint, a feedback loop exists between the nitrogenase function of bacteroids and the availability of arginine within the host plant. Ammonium released by bacteroids is rapidly incorporated by plant cells into glutamate, glutamine, and aspartate that all serve as precursors for the biosynthesis of ornithine and subsequently for arginine occurring within chloroplasts. The output of the CATCH-N cycle results in a net gain of assimilated nitrogen that subsequently amplifies the plant’s arginine biosynthesis capacity as part of a positive feedback mechanism. As humanity faces global challenges with population growth and climate change, we need to rethink how tomorrows agriculture will look like. Thereby systems-biology approaches to broaden our understanding of plant-microbe interactions, as well as the design of synthetic nitrogen-fixing microbes that mimic natural symbiosis with plants, hold significant promise. Our integrated model of the CATCH-N cycle provides new insights into the principles underlying legume symbiosis and comprises an important stepping stone for the rational biotechnological engineering of artificial nitrogen-fixing microbes and improved crop plants to ensure food and climate security.

## Materials and Methods

Supplementary Materials and Methods include detailed descriptions of bacterial strains and growth conditions, TnSeq library generation and data analysis, plant cultivation and phenotypic characterization, bacteroid extraction and substrate-specific ATP and nitrogenase stimulation, isotope tracing experiments, enzymatic procedures, and arginine catabolism thermodynamic calculations. Data SI contains the essentiality classification of each *S. meliloti* coding sequence across every selection screen.

### TnSeq library generation

Tn*5* hyper-saturated transposon mutant libraries in *S. meliloti* were generated as previously described (33, 35). Transposon mutant libraries were selected on rich medium (LB) supplemented with gentamicin and streptomycin. Plates were incubated at 30°C for 2 days. For the *in planta* selection, 4500 *Medicago truncatula* lss super-nodulator mutant (38) were flood-inoculated with an *S. meliloti* Tn*5* mutant reference libraries initially selected on rich media conditions (LB). Six weeks post-inoculation, nodules were retrieved, surface sterilized, crushed in cold PBS pH 7.4 and filtered through three layers of gauze to remove debris. The filtered suspension containing bacteroids was plated on LB supplemented with gentamicin and streptomycin and grown for 2 days at 30 °C. For all the selective conditions, the recovered *S. meliloti* colonies were pooled and stored at −80°C for subsequent use.

### Bacteroid isolation, substrate-specific stimulation of nitrogenase activity, and ATP production

*S. meliloti* and *B. diazoefficiens* bacteroids were isolated from *M. truncatula* (10 weeks post-inoculation) and *G. max* (3 weeks post-inoculation) root nodules respectively according to the following procedure. Under aerobic (ATP production stimulation) or anaerobic (for nitrogenase activity stimulation) conditions, nodules (0.25-1g fresh weight) were crushed in PBS pH=7.4 and filtered through three layers of gauze to remove debris. Under anaerobic conditions (for both assays), bacteroids were resuspended in 2 mL of induction media (2*μ*M biotin, 1 mM MgSO_4_, 42.2 mM Na_2_HPO_4_, 22 mM KH_2_PO_4_, 8.5 mM NaCl, 21 nM CoCl_2_, 1 *μ*M NaMoO4, pH 7.0) or nodule crude extract and added to 15 mL sealed flasks. Induction media was supplemented with either 7.4 mM succinate or 5 mM arginine or both substrates. For nitrogenase activity stimulation, acetylene and oxygen were added to a final concentration of 5 % and 0.01 % respectively in the head space of each flask. Nitrogenase activity was determined by the acetylene reduction assay. For ATP production, ATP content was determined for each sample through ATP-dependent luciferase reaction (BacTiter-Glo Microbial Cell Viability Assay, Promega) as indicated by the manufacturer.

### Enzymatic assays

Enzymatic reaction were done using purified C-terminal His 6X tag constructs (transaminases, urehydrolases, decarboxylases and *in vitro* synthetic reconstitution of arginine catabolism) or cell-lysates (dehydrogenases) from *E. coli* BL21 rosetta pLys strains expressing *S. meliloti* enzymes. Enzymatic reactions were done using 20-30 *μ*g of enzyme in 200 *μ*L reaction mixture containing 1 mM of substrate, 1 mM MgCl_2_, 1 mM MnCl_2_, 100 *μ*M piridoxal phosphate (for transaminase reactions), 1 mM NAD^+^ (for dehydrogenase reactions) and 500 *μ*M thiamine pyrophosphate (for decarboxylase reactions) in PBS. PBS pH was adjusted to 7.4 (transaminases), 8.0 (ureohydrolases, decarboxylases, *in vitro* synthetic reconstitution of arginine catabolism) or 10.0 (dehydrogenases). Enzymatic reactions were performed at 25 °C. For transaminases, urehydrolases, decarboxylases and *in vitro* synthetic reconstitution of arginine catabolism substrate consumption as well as product increase was detected by non-targeted *in vitro* metabolomics (52). Dehydrogenase reaction was determined by the conversion of NAD^+^ into NADH + H^+^, which was measured by the increase in absorbance at 340 nm.

## ACKNOWLEDGMENTS

We thank R. Schlapbach, L. Poveda from ZFGC for sequencing support, Anne Greet Bittermann and members from ScopeM, ETHZ for electron microscopy support, T. Fuhrer for metabolite analysis by LC-MS/MS. H. Christen for the conception of computational algorithms, mathematical conception and the statistical framework for TnSeq data analysis. R. Ledermann for the cultivation of *B. diazoefficiens*, H-M. Fischer for nitrogenase activity assay, isotope labeling experiments on isolated *B. diazoefficiens* bacteroids and comments on the manuscript. This work received institutional support from the Swiss Federal Institute of Technology (ETH) Zürich, ETH research grant [ETH-08 16-1] to B.C, the Swiss National Science Foundation, [31003A_166476, 310030_184664 and CRSII5_177164] to B.C and a Community Science Program (CSP) DNA sequencing award [CSP-1107] from the U.S. Department of Energy Joint Genome Institute in Walnut Creek. CA, USA. The work conducted by the U.S. Department of Energy Joint Genome Institute, a DOE Office of Science User Facility, is supported by the Office of Science of the U.S. Department of Energy under Contract No. DE-AC02-05CH11231.

