## Supplementary Information for "Co-catabolism of arginine and succinate drives symbiotic nitrogen fixation" (from another version of this preprint: version 1, 2019-08-21)

### This PDF file includes:

Supplementary Material and Methods
Figures S1 to S2
Tables S1 to S2
Supplementary References

**Bacterial strains, cultivation and growth conditions.** *Sinorhizobium meliloti* strain CL150 (Rm1021, *pstC+ ecfR1+*) (1) was grown at 30℃ in LB broth medium with 5 g NaCl per liter. *Escherichia coli* was grown in LB broth at 37℃. Where necessary, growth media were supplemented with antibiotics at the following concentrations: gentamicin, 10 µg/ml for *E. coli* and 30 µg/ml for *S. meliloti* when cultured in LB; streptomycin, 200 µg/ml; ampicillin, 50 µg/ml.

**Plant cultivation and inoculation assays.** *M. truncatula* WT Jaemalong seeds were surface sterilized with 70 % ethanol for five minutes and thoroughly rinsed with water. Seeds were imbibed with gentle agitation for at least four hours with two water changes and further imbibed overnight at room temperature without light. After imbibition, seeds were washed with water and germinated at 30℃ for 24 hours. Seedlings were planted into a sterile Perlite substrate (Isoself, Knauf) within 300 $cm^3$ black plastic pots (Greenhouse, Elho). Plants were grown in plant growth shelves at room temperature with a controlled 16 h day and 8 h night cycle. During the light cycle, every pot received 2500 lumens using 36 W Fluora 77 OSRAM light bulbs. Plants were automatically watered with a droplet watering system (Micro-drip-system, Gardena Art. 8311-20 Art. 1407-20) at three-day intervals with 80 mL of 10 % BNM solution (2). Pots were covered with a transparent PET cylinder during growth. Three days after germination, plants were inoculated with 20 mL of a *S. meliloti* cell culture with a cell density of an $OD_{600\ nm}$ of 0.05 resuspended in 1 mM $MgSO_4$. The inoculation time point was considered as day zero post-inoculation (dpi).

*G. max* cultivar Williams seeds were surface sterilized with a wash in 100 % ethanol for five minutes followed by a wash in 35 % $H_2O_2$ for 15 min. Afterward, seeds were thoroughly rinsed with water. Seeds were germinated in 0.8 % agar plates for 24 to 48 h at 25℃ in the dark. Seedlings were planted on 200 $cm^3$ brown glass jars filled with autoclaved vermiculite (Vermisol) and 100 mL of Jaensen media (3). After transplantation, plants were inoculated with *B. diazoefficiens* (1 mL $OD_{600\ nm}$ of 0.01). The inoculation time point was considered as day zero post-inoculation (dpi). Plants were grown in plant growth chambers at 28℃ with a controlled 16 h day and 8 h night cycle as previously described (4). After 5 dpi, plants were watered every 3 days with sterile water.

**Phenotypic characterization of plants to assess symbiotic nitrogen fixation.** Symbiosis phenotypes such as nodulation frequency, plant dry weight, and nitrogenase activity were determined in *M. truncatula* WT plants eight weeks post-inoculation with *S. meliloti* gene deletion mutants and wildtype control strains. Nitrogenase activity of *M. truncatula* was determined by measuring acetylene reduction to ethylene as previously reported. Plants were placed into 50 $cm^3$ sealed vials to which acetylene was injected to a final concentration of 2 %. Ethylene production was measured after 3 h and 5 h of incubation on an analytic gas chromatograph instrument (GC6850, Agilent Technologies). To determine the average ethylene production per nodule, ethylene production was normalized by nodule number and duration of acetylene incubation. Nodulation frequency was determined by manually counting all the visible nodules per plant. For dry weight determination, plants shoots were dried at 85℃ for at least 20 h prior individual plant weight measurement (ABS 80-40 N, KERN). Measurements from replicas were averaged and normalized to the *S. meliloti* CL150 WT reference strain. Nitrogenase activity, plant dry weight, and nodulation frequency were determined for at least 22 independent plants for each assayed *S. meliloti* strain.

***S. meliloti* bacteroid ultrastructure determination through scanning electron microscopy.** *M. truncatula* nodules were harvested 10 weeks post inoculation (wpi), cut open longitudinally and collected in a vessel with fixative buffer (2.5 % Glutaraldehyde, 2 % Formaldehyde in 0.15 M Na-Cacodylate with 2 mM $CaCl_2$). The nodule containing vessel was subjected vacuum to for at least 30 min to allow the nodules to sink followed by microwave fixation in fresh fixative (Pelco Biowave Pro+) and washed with fixative buffer. Nodules were stained through sequential incubation with (*1*) 2 % $OsO_4$, 1.5 % $K_4Fe(CN)_6$ in 0.15 M Na-Cacodylate with 2 mM $CaCl_2$; (*2*) 1 % Thiocarbohydrazide; (*3*) 2 % $OsO_4$; (*4*) 1 % Uranylacetate; (*5*) Waltons lead aspartate; samples were washed with fixative buffer between every step. Samples were dehydrated through a graded ethanol series (25 %, 50 %, 75 %, 100 %, dry ethanol) and washed with dry acetone. Samples were placed in a graded series of Epon/Araldite in dry acetone (25 %, 50 %, 75 %, 100 %, 100 %) and allowed to polymerize at 60℃ for three days. Ultrathin sections (100 nm) were transferred to Si-wafer-chips and imaged on a FE-SEM FEI Magellan 400i operating at 1,8 kV and 0,8 nA with a 20 nm pixel size by backscatter electron detection.

**Substrate specific stimulation of nitrogenase activity in isolated bacteroids.** *S. meliloti* and *B. diazoefficiens* bacteroids were isolated from *M. truncatula* and *G. max* root nodules respectively according to following procedure. Plant roots were harvested 3 weeks (*G. max*) or 10 weeks (*M. truncatula*) post inoculation with wild-type *B. diazoefficiens* 110spc4 strain and *S. meliloti* CL150 strain respectively and rinsed thoroughly with water. Under anaerobic conditions (92 % $N_2$, 8 % $H_2$), nodules (250 mg to 1 g wet weight) were crushed in PBS (10 mM $Na_2HPO_4$, 1.8 mM $KH_2PO_4$, 137 mM NaCl, 2.7 mM KCl, pH 7.4) and filtered through three layers of gauze to remove debris. Bacteroids suspensions (2 mL corresponding to 4 * $10^6$ bacteroid cells) were added to 15 mL sealed flasks to obtain reference measurements of nitrogenase activity in isolated bacteroids in presence of plant crude extract. To assay stimulation of nitrogenase activities upon addition of substrates, bacteroid suspensions were washed two times with PBS under anerobic conditions and pelleted by centrifugation at 1000 *x g*. Recovered bacteroids (4 * $10^6$ cells) were resuspended in 2 mL of induction media (2 $\mu$M biotin, 1 mM $MgSO_4$, 42.2 mM $Na_2HPO_4$, 22 mM $KH_2PO_4$, 8.5 mM NaCl, 21 nM $CoCl_2$, 1 $\mu$M $NaMoO_4$ pH 7.0) and added to 15 mL sealed flasks. Induction media was supplemented with either 7.4 mM succinate or 5 mM arginine or both substrates. Acetylene and oxygen were added to a final concentration of 5 % and 0.01 % respectively in the head space of each flask. Nitrogenase activity was determined for each sample through measuring the reduction of acetylene into ethylene after 1 hour and 3 hour time points. Ethylene production was detected with gas chromatograph (GC6850, Agilent Technologies).

**Substrate specific stimulation of ATP production in isolated bacteroids.** *B. diazoefficiens* bacteroids were isolated from 3 week post inoculated *G. max* root nodules. Nodules (1g wet weight) were crushed in PBS (10 mM $Na_2HPO_4$, 1.8 mM $KH_2PO_4$, 137 mM NaCl, 2.7 mM KCl, pH 7.4) and filtered through three layers of gauze to remove debris. Bacteroid suspension was pelleted by centrifugation at 2500 *x g* and the supernatant (nodule extract) was saved for later usage. Bacteroid suspension (5 * $10^8$ cells) was washed twice with PBS and resuspended in 1 mL of induction media (2 $\mu$M biotin, 1 mM $MgSO_4$, 42.2 mM $Na_2HPO_4$, 22 mM $KH_2PO_4$, 8.5 mM NaCl, 21 nM $CoCl_2$, 1 $\mu$M $NaMoO_4$ pH 7.0). To avoid ATP generation from aerobic respiration, the bacteroid suspension was placed under anaerobic conditions (92 % $N_2$, 8 % $H_2$) to perform the rest of the procedure. Bacteroids (200 $\mu$L) were incubated in 2 mL of nodule extract (obtained at the beginning of the protocol) or induction media without supplements or supplemented with either 7.4 mM succinate or 5 mM arginine or both substrates. ATP content was determined for each sample (using a 1:10 dilution) through ATP-dependent luciferase reaction (BacTiter-Glo Microbial Cell Viability Assay, Promega). Luminiscence from luciferase activity was quantified with Victor3 multilabel plate counter (PerkinElmer).

**Transposon mutagenesis.** To generate hyper-saturated transposon mutant libraries in *S. meliloti*, a previously described Tn5 mutagenesis procedure for *Caulobacter crescentus* was adapted (5). In brief, the Tn*5* delivery ColE1 plasmid pTn*5*_gent_14N (6) was conjugated from an *E. coli* SM10 donor strain into a *S. meliloti* CL150 recipient strain. Separate transposon mutant libraries were generated and growth selected on rich medium (LB). For each condition, a total of sixteen independent conjugations were performed and replicate libraries were tagged using eight barcoded Tn*5* derivatives. Transposon insertion mutants were selected on LB supplemented with gentamicin and streptomycin. Plates were incubated at 30℃ for 2 days and transposon mutant libraries from each plate were separately pooled, supplemented with 10 % v/v DMSO (Sigma Aldrich), and stored in 96 well deep well plates at -80℃ for further processing.

***In planta* selection of transposon mutant libraries.** Transposon mutant pools were selected for infection and nodule formation in legume plants. For the nodulation experiment, a *Medicago truncatula* lss super-nodulator mutant was used (7). Seeds were treated with concentrated $H_2SO_4$ (Sigma-Aldrich) for 5 min, thoroughly rinsed with sterile water, the sterilized with 7 % NaClO (VWR chemicals) for 3 min and again rinsed with sterile water. The seeds were then imbibed with gentle agitation for four hours with regular water changes and then incubated overnight in the dark at room temperature. After imbibition, seeds were rinsed with sterile water, placed in deep petri plates and inverted for 24 hours at 30 ℃ to allow for the downward growth of the seedling roots. After removing seed coats, groups of 25 seedlings were planted on large square plates (Genetix) containing 1.2 % buffered nodulation medium (2) supplemented with 0.1 nM aminoethoxy vinyl glycine. Altogether, 4500 *M. truncatula* lss plant seedlings were grown at 22 ℃ with a controlled 16h day and 8h night cycle (2500 Lumens using Osram Fluora L36 W/77 bulbs). Five days post-germination, *M. truncatula* lss seedlings were flood-inoculated with *S. meliloti* Tn5 mutant reference libraries initially selected on rich media conditions (LB). During inoculation experiments, input libraries were kept independent from each other. *S. meliloti* transposon input mutant pools were inoculated from 96 well storage plates, grown overnight, washed, and resuspended to an $OD_{600\ nm}$=0.05 in 10 mM $MgSO_4$. Plant roots were then aseptically inoculated with 5 mL of a diluted bacterial suspension, followed by removal of the excess bacterial suspension.

**Recovery of transposon mutant libraries from nodules.** After six weeks post-inoculation, nodules were harvested. To recover transposon mutants capable of infecting root nodules, a two-step surface sterilization protocol was employed. Nodule material was washed with 1 % SDS, then treated with 70 % ethanol for 5 min, followed by rinsing with sterile water. Next, nodules were treated for 3 min with 0.2 % NaClO (VWR chemicals) followed by three rinses with sterile water. The surface-sterilized nodules were crushed in cold PBS (10 mM $Na_2HPO_4$, 1.8 mM $KH_2PO_4$, 137 mM NaCl, 2.7 mM KCl, pH 7.4) and filtered through three layers of gauze to remove debris. The filtered suspension containing bacteroids was plated on LB supplemented with gentamicin and streptomycin and grown for 2 days at 30℃. The recovered *S. meliloti* colonies were pooled and arrayed in 96 well plates, and stored at -80ºC for subsequent use.

**Data analysis and mapping of transposon insertion sites.** Raw sequencing data processing and read alignment were performed using a custom sequence analysis pipeline based on Python, Biopython (8), bwa (9), and Matlab routines as previously described (6). Adapter sequences were detected using Python string comparison with a 15 bp search window. Demultiplexing into the different TnSeq selection experiments was performed according to a defined barcode sequence tag internal to the arbitrary primer sequence. Reads were aligned onto the *Sinorhizobium meliloti* 1021 NCBI reference genome (NC_003047, NC_003037, NC_003078 (10–12)) using bwa-07.12 (9). Insertion data sets were correlated with the genome annotation to analyze global insertion statistics and calculate transposon insertion occurrence and distributions within each annotated GenBank feature of the *S. meliloti* 1021 genome (10) as previously described (6).

**Gene essentiality analysis across selection conditions.** . The metrics and statistical analysis for classification of protein-coding sequences (CDS) into essential, promoting fitness, and non-essential categories have been previously described (5). CDS are classified as "essential" or "fitness" genes according to the following criteria. CDS are classified as essential if the transposon insertion density is less than six times the average insertion density measured across the genome and has either a single transposon insertion gap covering more than 60 % of the CDS or has two internal transposon gaps covering more than 80 % of the CDS. If the average transposon density is less than four times the average insertion density across the genome but no essentiality criterion was satisfied, CDS were classified as fitness genes. For the nodule selected TnSeq dataset, a symbiosis impairment score (*p*-values) was calculated for each gene based on the number and distribution of transposon insertions recovered from the *in planta* selection experiments. The symbiosis impairment score was calculated for each gene as the product of the probability *p1*, to recover *i* transposon insertions when $\lambda$ insertions are expected according to a Poisson distribution, and the probability *p2*, to observe a consecutive loss in transposon insertions equals or exceeding *k* insertions as compared to the input mutant pool. The expected number of recovered insertions was estimated based on the observed loss-rate of neutral insertions ($l = 6.840$) due to sampling. The probability *p1* was calculated according to the following formula:

$$p1 = \sum_{s=0}^{i} \frac{(e^{-\lambda}) \cdot (\lambda^{s})}{s!}$$

The probability *p2* was calculated for each gene based on the total number of insertions (*n*) present in the input mutant pool, the number of recovered insertions from the output mutant pool (*i*) and the maximal number of consecutive insertions that where lost upon *in planta* selection (*k*) by calculating the number (h) of compositions of length *i+1* for *n* where each part does not exceed *k*, divided by the total number of compositions ($h_{tot}$) for *n*. The probability *p2* was calculated according to the following formula.

$$h = \frac{(i+1)\cdot(n-k)!}{(i)!\cdot(n-i-k)!} \qquad h_{tot} = \frac{n!}{(i)!\cdot(n-i)!} \qquad p_2 = \frac{h}{h_{tot}} \qquad \text{for } k > \frac{n-i}{2}$$

For *k* smaller than (n-i)/2 and *n* not to exceed 100, *p2* was numerically calculated by calculating the number (h) of compositions of length *i+1* for *n* where each part does not exceed *k* by brute force. For *k* smaller than (n-i)/2 and *n* equal or exceed 100, we approximated p2 by sampling the composition space by random simulation and counting the occurrences of compositions of length *i+1* for *n* where each part does not exceed *k* .

**Construction of targeted gene deletions in *Sinorhizoium meliloti*.** *S. meliloti* deletion mutants were generated by replacing the native gene with the *aacC1* gene conferring gentamycin resistance using a one-step double homologous recombination procedure as detailed in (13). Flanking DNA regions covering 750 bp upstream and downstream of a target gene were PCR amplified and subsequently fused to a central gentamicin resistance gene using splicing by overlapping extension PCR or Gibson assembly (14) to produce gene replacement cassettes. The gene replacement cassettes were cloned via SpeI and MfeI into the pNPTS138 plasmid, which is non-replicative in *S. meliloti* and confers kanamycin resistance. Cloned plasmids were sequence confirmed and conjugated into *S. meliloti* strain CL150. Recombinants were selected on LB media supplemented with gentamicin and subsequently screened for kanamycin sensitivity. Single gene deletion strains were confirmed by PCR and sequencing. A similar deletion strategy was employed for construction of the double arginase mutant ($\Delta$*argI1* $\Delta$*argI2*) with a gene replacement cassette for *argI1* carrying a spectinomycin marker gene. The resulting plasmid was conjugated into *S. meliloti* CL150 $\Delta$*argI2* and double arginase double mutant strain were screen for kanamycin sensitivity prior verification of constructed deletion by PCR and sequencing.

**$^{13}$C Arginine and unlabeled succinate feeding in *B. diazoefficiens* bacteroids.** *B. diazoefficiens* bacteroids were isolated under anaerobic conditions from three-week post-inoculated *G. max* root nodules according to (15, 16). Nodules (10 g wet weight) were crushed in PBS (10 mM $Na_2HPO_4$, 1.8 mM $KH_2PO_4$, 137 mM NaCl, 2.7 mM KCl, pH 7.4). The homogenate was passed through four layers of cheesecloth (pre-moistened with PBS) into a sterile centrifuge tube, re-extracted several times with buffer and centrifuged at 400 *x g* for 10 min. The resulting pellet was re-suspended twice in PBS and centrifuged at 8000 *x g* for 20 min. The pellet was dispersed into the extraction buffer (2 mL/g original weight of the nodule) and was layered onto a pre-equilibrated Ludox gradient consisting of 25 % (10 mL) Ludox, and 75 % PBS (30 mL) The gradient tubes were centrifuged in a SW-28 rotor at 10 000 *x g* for 35 min at 4℃ in a Beckman L8–55 ultracentrifuge. The bacteroid layer with a density of 1.09 g/ml was collected. The bacteroid pellet was suspended in distilled water and collected by centrifugation at 10000 g. For labeling assays, bacteroids were resuspended in 10 ml Volume in PBS to a final OD of 4.0. After the addition of succinate (5 mM) and $^{13}$C arginine, bacteroids were incubated at RT under microaerobic condition (0.1 % v/v) and samples of 250 µl were taken at regular time intervals, filtered on a PVDF 0.45 µl membrane and immediately washed with 1.0 ml $H_2O$ chromasolv, the filter with the cells was placed in 3 ml (40 % MeOH, 40 % Formamide and 20 % $H_2O$) for extraction of metabolites, storage at -80℃, lyophilized in a speedvac.

**Expression and purification of recombinant proteins.** Coding sequences of interest were amplified by PCR and cloned into pET42 expression plasmids inframe to a C-terminal (His)6-tag (6xHis). The resulting vectors were sequence-verified and electroporated into BL21 rosetta pLys strains. *E. coli* BL21 harboring the expression vectors were grown at 30℃ in LB medium containing chloramphenicol (20 mg/liter) and kanamycin (30 mg/liter). When the cultures reached an $OD_{600nm}$ of 0.4, isopropyl-$\beta$-D-thiogalactopyranoside (IPTG) was added to a final concentration of 0.5 M. After the addition of IPTG, the culture was grown for 2-4h more at 30℃ to induce the expression of the recombinant proteins. Grown cells were harvested by centrifugation at 5095 *x g* for 10 min at 4℃. The resulting pellet was either shock frozen with liquid $N_2$ and stored at -80℃ or immediately used for protein extraction and purification. To purify the recombinant proteins, cells were resuspended in NPI-buffer (20 mM sodium phosphate, 500 mM NaCl, pH7.4) supplemented with 10 mM imidazole and disrupted using a French press (SLM Instruments Inc.). Lysates were cleared by centrifugation at 17000 x g for 15 min at 4°C to remove cell debris. The supernatant was loaded on HisTrap FF crude column (GE Healthcare) previously equilibrated with NPI-buffer supplemented with 10 mM imidazole. The column was washed twice with NPI-buffer supplemented with 10 mM and 20 mM imidazole respectively. The purified enzymes were eluted with NPI-buffer supplemented with 250 mM imidazole, concentrated by ultrafiltration with Amicon Ultra-4 centrifugal filters (Merck Millipore) and dialyzed against NPI-buffer containing 10 % (v/v) glycerol. Protein concentration was determined using Pierce BCA Protein Assay kit (Thermo Scientific). Protein samples were shocked frozen with liquid $N_2$ and stored at -80°C until use.

**Chemical synthesis of succinate semialdehyde, 4-aminobutanal and 4-guanidinobutanal.** The arginine transamination metabolic network contains a set of 13 intermediates. Thereof, three intermediates succinate semialdehyde, 4-aminobutanal and 4-guanidinobutanal are not commercially available and were chemically synthesized for use in subsequent enzyme characterization studies as aldehyde dehydrogenase substrates. Succinate semialdehyde was synthesized according to the procedure as previously described (17). In a 15 mL Falcon tube, a solution of monosodium glutamate (169 mg, 1 mmol) in 5.0 mL distilled water was exposed to a gentle flow of $N_2$ for 5 min. An equimolar amount of chloramine T (227 mg, 1 mmol) was added to the solution and dissolved by heating the solution to 60°C. After further incubation at 60°C for an additional 15 min the mixture was cooled to 25°C and adjusted to pH 2.0 with concentrated HCl and degassed. Reaction by-products were crystallized by placing the mixture on ice and removed by filtration. The aqueous phase was extracted with diethyl ether 3 times, organic phases where combined and the water fraction was discarded. Diethyl ether was evaporated yielding an aqueous solution of 1 % of the starting material and a concentration of approximately 2 M succinic semialdehyde. 4-guanidinobutyraldehyde was prepared from L-arginine according to (18). The procedure follows a similar reaction scheme as detailed for the synthesis of succinic semialdehyde with the following modifications. As starting material, arginine hydrochloride (210 mg, 1 mmol) was dissolved. Upon addition of chloramine T (227 mg, 1 mmol), the mixture was adjusted to a pH 6.5 with 1 N HCl (50 $\mu$l) and heated to 60°C. Prior extraction with diethyl ether, the solution was adjusted to pH 13.5 with 10 N NaOH. 4-aminobutyraldehyde was prepared by the hydrolysis of 1.0 mL of 0.5 M 4-aminobutyraldehyde diethyl acetal (Sigma-Aldrich). 172 $\mu$l 4-aminobutyraldehyde diethyl acetal was dissolved in 2 mL $H_2O$ and 1 N HCl was added to acidify to pH 3.0. After incubation for 30 min at 30°C, the reaction mixture was titrated to pH 10.0 with 1 N NaOH. The crude reaction product 4-aminobutanal was extracted from the water phase with five consecutive extraction steps using 1 mL diethyl ether. Ether fraction were combined and solvent was removed by evaporation. The reaction product 4-aminobutyraldehyde was obtained as colourless liquid of 40.0 mg mass.

**Transaminase *in vitro* reaction assay.** Transaminases were purified as C-terminal His 6X tag constructs from *E. coli* BL21 Rosetta pLys strains and assayed for enzymatic activity according to following procedure. A total of 30 $\mu$g purified enzyme were added to 200 $\mu$L reaction mixture containing 1 mM arginine, 1 mM ornithine, 1 mM citrulline, 1 mM agmatine, 1 mM putrescine, 1 mM 4-guanidinobutanoate, 1 mM 4-aminobutanoate, 10 mM of pyruvate, 1 mM $MgCl_2$, 1 mM $MnCl_2$, 100 μM piridoxal phosphate in 50 mM PBS pH 7.4 followed by incubation at 25°C. Aliquots (20 $\mu$L) of the enzymatic reaction were sampled over the time series and reaction were blocked by adding 180 $\mu$l of ice-cold methanol. Substrate consumption as well as product increase was detected by non-targeted *in vitro* metabolomics (19).

**Ureohydrolase *in vitro* reactions assay.** Ureohydrolases were purified as C-terminal His 6X tag constructs from *E. coli* BL21 Rosetta pLys strains and assayed for enzymatic activity according to the following procedure. A total of 30 $\mu$g purified enzyme were added to 200 $\mu$L reaction mixture containing 1 mM arginine, 1 mM agmatine, 1 mM 4-guanidinobutanoate, 10 mM of pyruvate, 1 mM $MgCl_2$, 1 mM $MnCl_2$, 100 $\mu$M pyridoxal phosphate in 50 mM PBS pH 8.0, previously incubated at 25°C for 30 min with 0.15 ng/ $\mu$L purified AspC followed by heat inactivation at 65°C for 5 min to generate 5-guanidino-2-oxopentanoate (GOP) and guanidinobutanal, from arginine and agmatine transamination respectively. Ureohydrolase reactions were done at 25°C. Aliquots (20 $\mu$l) of the enzymatic reaction were sampled over the time series and reaction was blocked by adding 180 $\mu$l of ice-cold methanol. Substrate consumption as well as product increase was detected by non-targeted *in vitro* metabolomics (19).

**Decarboxylase *in vitro* reactions assay.** Decarboxylases were purified as C-terminal His 6X tag constructs from *E. coli* BL21 Rosetta pLys strains and assayed for enzymatic activity according to following procedure. A total of 30 $\mu$g purified enzyme were added to 200 $\mu$L reaction mixture containing 1 mM arginine, 1 mM ornithine, 1 mM citrulline, 10 mM of pyruvate, 1 mM $MgCl_2$, 1 mM $MnCl_2$, 100 $\mu$M piridoxal phosphate, 500 $\mu$M thiamine pyrophosphate in 50 mM PBS pH 8.0, previously incubated at 25°C for 30 min with 0.15ng/ $\mu$L purified AspC followed by heat inactivation at 65°C for 5 min to generate 5-guanidino-2-oxopentanoate (GOP) from arginine transamination. Decarboxylase reactions were done at 25°C. Aliquots (20 $\mu$l) of the enzymatic reaction were sampled over the time series and reaction were blocked by adding 180 $\mu$l of ice-cold methanol. Substrate consumption as well as product increase was detected by non-targeted *in vitro* metabolomics (19).

**Dehydrogenase *in vitro* reactions assay.** Dehydrogenase activities were assayed from cell-lysates of BL21 rosetta pLys strain expressing *S. meliloti* dehydrogenases according to the following procedure. Cell-lysates corresponding to 20 $\mu$g of dehydrogenase enzymes were added to a substrate mixture containing 1 mM succinate semialdehyde or 1 mM 4-Guanidinobutanal or 1 mM 4-aminobutanal in 10 mM PBS pH 10.0 supplemented with 1 mM $NAD^+$. Dehydrogenase reaction was determined by the conversion of $NAD^+$ into NADH + $H^+$, which was measured by the increase in absorbance at 340 nm.

**Biochemical reconstitution of the catabolic arginine transamination network.** To reconstruct a functional catabolic arginine transamination network *in vitro*, a multienzyme assay comprising 14 purified enzymes was established. A reaction buffer containing of 500 $\mu$M thiamine pyrophosphate, 100 $\mu$M pyridoxal phosphate, 2 mM NAD, 1 mM $MgCl_2$ and 1 mM $MnCl_2$ in 50 mM PBS pH 8.0 was prepared. Purified enzymes (25 $\mu$g each) from the catabolic arginine transamination network (AspC, AatB, ArgD, GabT2, DatA, ArgI1, SpeB, SpeB2, IlvB1, OdcA, OdcB, GabD1, GabD6, and GabD7) were added one by one and gently mixed into the reaction mixture. After the addition of all enzymes, 20 mM pyruvate and 2 mM arginine were added and gently mixed. As a control, the enzyme mix was incubated in a reaction buffer lacking arginine and pyruvate. Aliquots (15 $\mu$l) of the enzymatic reaction were sampled over the time series and reaction was blocked by adding 135 $\mu$l of ice-cold methanol. Substrate consumption as well as product increase was detected by non-targeted *in vitro* metabolomics (19).

### Supplementary Figures

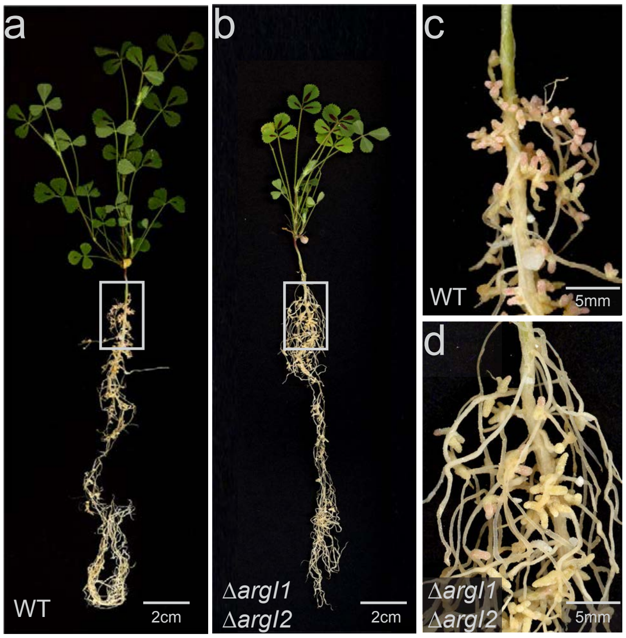


**Figure S1.** *Symbiotic defects of S.meliloti mutants impaired in arginine catabolism*. (**a**,**b**) Assessment of nitrogen starvation phenotype of *M. truncatula lss* plants inoculated with*S. meliloti* WT (a) or the *S. meliloti double deletion mutant* Δ*argl1*,Δ*argl2.* (**c**,**d**) Close up image of the root systems from the plants shown in ***a*** and ***b*** respectively.

S. *meliloti* bacteroid ultrastructure

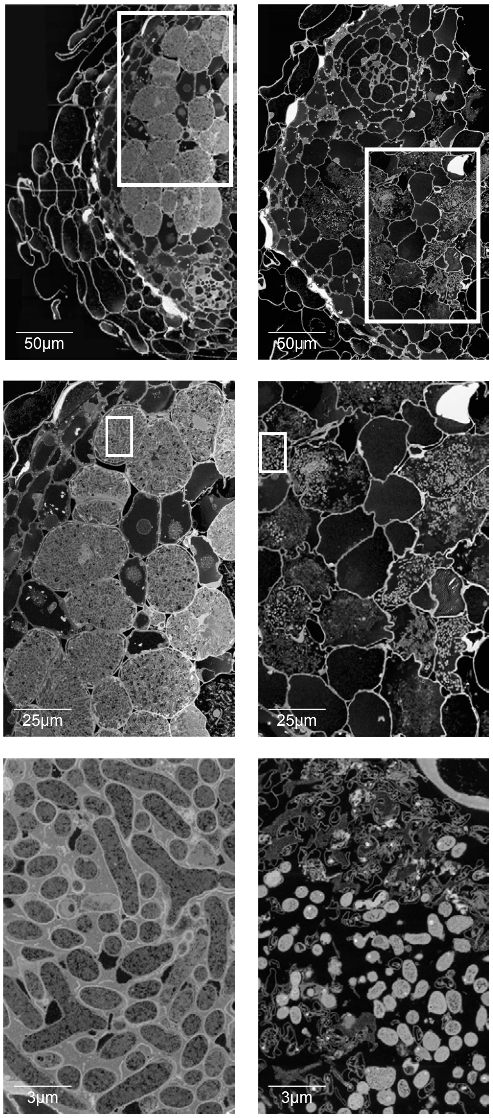


**Figure S2.** Arginine catabolism is fundamental for bacteroid development. Scanning electron micrographs from cross-sections of nodules bearing *S. meliloti* WT (left) or arginine catabolism double mutant △ *argI1* △ *argI2* (right) showing bacteroid occupancy of the nodule and ultrastructure. The white box indicates the area enlarged for the subsequent panel.

### Supplementary Tables

**Table S1. Symbiotic $N_2$-fixation phenotypes of *M. truncatula* inoculated with *S. meliloti***

| *S. meliloti* | Shoot weight[a] | Nodulation frequency[b] | Nitrogenase activity [%][c] | Replicates |
|---|---|---|---|---|
| WT | 18.15 ± 1.42 | 17.95 ± 0.75 | 100.00 ± 7.36 | 109 |
| Δ*artABCDE* | 15.59 ± 1.73 | 28.39 ± 1.70 | 52.94 ± 7.27 | 46 |
| Δ*satABC* | 12.81 ± 1.89 | 25.58 ± 2.25 | 44.55 ± 10.99 | 33 |
| Δ*argI1* | 10.51 ± 0.93 | 27.06 ± 2.28 | 28.82 ± 5.21 | 47 |
| Δ*argI2* | 11.34 ± 1.20 | 26.84 ± 1.72 | 29.03 ± 5.16 | 43 |
| Δ*argI1*Δ*argI2* | 9.79 ± 1.11 | 36.70 ± 2.57 | 19.11 ± 3.15 | 40 |
| Δ*ureGFE* | 9.46 ± 0.62 | 26.43 ± 2.27 | 19.10 ± 4.81 | 30 |
| Δ*amtB* | 12.73 ± 1.25 | 30.97 ± 2.37 | 35.32 ± 5.50 | 31 |
| Δ*aspC* | 15.45 ± 1.65 | 27.75 ± 1.54 | 49.09 ± 9.00 | 40 |
| Δ*dctAB* | 7.01 ± 0.29 | 30.77 ± 2.14 | 0.00 ± 0.04 | 35 |
| Δ*nifD* | 6.60 ± 0.23 | 35.49 ± 1.67 | 0.00 ± 0.03 | 72 |
| Non inoculated | 6.31 ± 0.31 | 0.00 ± 0.00 | n.a.[d] | 42 |

[a] Shoot mass of *M. truncatula* plants after 8 week post inoculation. Shown is average ± SEM.
[b] Number of visible nodules per plant after 8 week post inoculation. Shown is mean ± SEM.
[c] Relative nitrogenase activity per nodule. Shown is mean ± SEM.
[d] Not applies as non inoculated controls did not form nodules.

**Table S2. Substrate specificity of the transaminase AspC**

| Amino donor | Relative activity (%)[a] |
|---|---|
| *L*-Arginine | 100.0 ± 4.9 |
| Agmatine | 112.6 ± 6.8 |
| Putrescine | 73.8 ± 6.8 |
| Ornithine | 64.6 ± 6.8 |

[a] The pyruvate transamination activity of AspC with *L*-arginine as the substrate was defined as 100%. Shown is the mean and standard error of the mean (SEM) of at least 4 independent replicates.
