## Supplementary material for "Co-catabolism of arginine and succinate drives symbiotic nitrogen fixation": Model_CATCH_N1 (from another version of this preprint: version 1, 2019-08-21)

**Reaction stoichiometry of nitrogen fixation metabolism upon co-feeding of arginine and succinate.** . Reaction equations are step-wise deduced from the conversion of arginine into gamma-amino butanoic acid (GABA), GABA to Succinate and conversion to Pyruvate. Furthermore, reaction equations are deduced from electron bifurcation of NADH by the FixABCX complex, nitrogen fixation by nitrogenase complex and oxidative phosphorylation by the FbcBCF and the high-affine terminal cbb3 oxidases FixNOPQ1-3.

**CATCH-N1, stochiometric model with pyruvate as the final amino-acceptor**
Arginine to GABA:

$$\mathrm{Arg} + \mathrm{NAD} + \mathrm{Pyr} + 3\,\mathrm{H_2O} \longrightarrow \mathrm{GABA} + \mathrm{Ala} + \mathrm{NADH_2} + 2\,\mathrm{NH_3} + 2\,\mathrm{CO_2}, \quad [1]$$

1 ATP generated and 3 $H^+$ consumed per arginine through the carbamoyl pathway [2]

GABA to succinate:

$$\mathrm{GABA} + \mathrm{Pyr} + \mathrm{NAD} + \mathrm{H_2O} \longrightarrow \mathrm{Suc} + \mathrm{Ala} + \mathrm{NADH_2} \quad [3]$$

Arginine to succinate, 1 + 3:

$$\mathrm{Arg} + 2\,\mathrm{NAD} + 2\,\mathrm{Pyr} + 4\,\mathrm{H_2O} \longrightarrow \mathrm{Suc} + 2\,\mathrm{Ala} + 2\,\mathrm{NADH_2} + 2\,\mathrm{NH_3} + 2\,\mathrm{CO_2} \quad [4]$$

Succinate to pyruvate:

$$2\,\mathrm{Suc} + 2\,\mathrm{Q} + 2\,\mathrm{NAD} + 2\,\mathrm{H_2O} \longrightarrow 2\,\mathrm{Pyr} + 2\,\mathrm{NADH_2} + 2\,\mathrm{QH_2} + 2\,\mathrm{CO_2} \quad [5]$$

Co-catabolism of arginine and succinate to 2 alanine: 4 + 5:

$$\mathrm{Arg} + \mathrm{Suc} + 4\,\mathrm{NAD} + 2\,\mathrm{Q} + 6\,\mathrm{H_2O} \longrightarrow 2\,\mathrm{Ala} + 4\,\mathrm{NADH_2} + 2\,\mathrm{QH_2} + 2\,\mathrm{NH_3} + 4\,\mathrm{CO_2} \quad [6]$$

Electron bifurcation of NADH by the FixABCX complex:

$$8\,\mathrm{NADH_2} + 8\,\mathrm{Fd_{ox}} + 4\,\mathrm{Q} \longrightarrow 8\,\mathrm{NAD} + 8\,\mathrm{Fd_{red}} + 4\,\mathrm{QH_2} \quad [7]$$

Nitrogenase reaction by NifDK (as biochemical equation not balanced for $H^+$):

$$\mathrm{N_2} + 8\,\mathrm{Fd_{red}} \longrightarrow 2\,\mathrm{NH_3(aq)} + \mathrm{H_2} + 8\,\mathrm{Fd_{ox}}, \quad \text{requires } 8\,\mathrm{H^+},\ 16\ \mathrm{ATP} \quad [8]$$

Oxidative phosphorylation by FbcBCF and high-affine terminal oxidase FixNOPQ and:

$$8\,\mathrm{QH_2} + 4\,\mathrm{O_2} \longrightarrow 8\,\mathrm{Q} + 8\,\mathrm{H_2O} \quad (8 \cdot 6\,\mathrm{H^+} = 48\,\mathrm{H^+}\ \text{exported}) \quad [9]$$

Overall biochemical reaction, not yet balanced for charge and $H^+$:

$$2\,\mathrm{Arg} + 2\,\mathrm{Suc} + \mathrm{N_2} + 4\,\mathrm{O_2} + 4\,\mathrm{H_2O} \longrightarrow 4\,\mathrm{Ala} + 6\,\mathrm{NH_3} + 8\,\mathrm{CO_2} + \mathrm{H_2} \quad \text{biochem. eq.} \quad [10]$$

Balancing the reaction stoichiometry for charge and $H^+$:

$$2\,\mathrm{C_6H_{15}N_4O_2}^{+} + 2\,\mathrm{C_4H_4O_4}^{2-} + \mathrm{N_2} + 4\,\mathrm{O_2} + 4\,\mathrm{H_2O} \longrightarrow 4\,\mathrm{C_3H_7NO_2} + 6\,\mathrm{NH_4}^{+} + 8\,\mathrm{CO_2} + \mathrm{H_2} \quad \text{chemical eq.} \quad [11]$$

$$20\,\mathrm{C} = 20\,\mathrm{C} \quad \text{C balanced} \quad [12]$$

$$10\,\mathrm{N} = 10\,\mathrm{N} \quad \text{N balanced} \quad [13]$$

$$24\,\mathrm{O} = 24\,\mathrm{O} \quad \text{O balanced} \quad [14]$$

$$46\,\mathrm{H} \neq 54\,\mathrm{H} \quad 8\,\mathrm{H^+}\ \text{missing} \quad [15]$$

$$-2 \neq +6 \quad +\,8\ \text{missing} \quad [16]$$

Insertion of 8 $H^+$ to the left of 11 results in the following charge and $H^+$ corrected equation:

$$2\,\mathrm{C_6H_{15}N_4O_2}^{+} + 2\,\mathrm{C_4H_4O_4}^{2-} + \mathrm{N_2} + 4\,\mathrm{O_2} + 4\,\mathrm{H_2O} + 8\,\mathrm{H^+} \longrightarrow 4\,\mathrm{C_3H_7NO_2} + 6\,\mathrm{NH_4}^{+} + 8\,\mathrm{CO_2} + \mathrm{H_2} \quad \text{chemical eq.} \quad [17]$$

Proton translocation:

$$48\,\mathrm{H^+}_{\mathrm{in}} \longrightarrow 48\,\mathrm{H^+}_{\mathrm{out}} \quad \mathrm{O_2}\ \text{dependent proton translocation from 9} \quad [18]$$

$$48\,\mathrm{H^+}_{\mathrm{out}} \longrightarrow 48\,\mathrm{H^+}_{\mathrm{in}} \quad \text{protons available from 9} \quad [19]$$

$$8\,\mathrm{H^+}_{\mathrm{out}} \longrightarrow 8\,\mathrm{H^+}_{\mathrm{in}} \quad \text{influx of } \mathrm{H^+} \text{ derived from 17} \quad [20]$$

$$56\,\mathrm{H^+}_{\mathrm{out}} \longrightarrow 56\,\mathrm{H^+}_{\mathrm{in}} \quad \text{19 + 20 } \textbf{Sum of proton influx} \quad [21]$$

Stochiometry of the proton motive force (PMF) dependent ATP synthesis:

$$\frac{[H^+]}{[ATP]} = \frac{10}{3} \quad , \frac{56}{3.33} = 16.8, \text{ ATP synthesized} \tag{22}$$

Stoichiometry of Enzyme-coupled ATP synthesis:

2 ATP/ 2 arginine generated by the action of deiminase, transcarbamoylase and carbamate kinase, 1 [23]

Sum of oxygen-dependent, oxygen-independent and Enzyme-coupled ATP synthesis - :

14.40 ATP Oxygen-dependent [24]
2.40 ATP Oxygen-independent [25]
2 ATP Enzyme-coupled [26]
18.8 ATP Sum [27]
-16 ATP Nitrogenase reaction [28]
**2.8 ATP Net gain** [29]
[30]

Overall reaction of $N_2$ fixation by bacteroid:

$$2\,\mathrm{Arg} + 2\,\mathrm{Suc} + \mathrm{N_2} + 4\,\mathrm{O_2} + n\,\mathrm{ADP} + n\,\mathrm{Pi} + (4\text{-n})\,\mathrm{H_2O} + 8\,\mathrm{H^+} \longrightarrow 4\,\mathrm{Ala} + 6\,\mathrm{NH_4}^+ + 8\,\mathrm{CO_2} + \mathrm{H_2} + \mathrm{nATP} \tag{31}$$

with n= 2.80 [32]
