## Supplementary material for "Co-catabolism of arginine and succinate drives symbiotic nitrogen fixation": Model_TCA (from another version of this preprint: version 1, 2019-08-21)

**Reaction stoichiometry of $N_2$ fixation metabolism under the assumption of an operational TCA cycle.** . Reaction equations are step-wise deduced from the conversion of malate and succinate respectively. Furthermore, reaction equations are deduced from electron bifurcation of NADH by the FixABCD complex, nitrogen fixation by nitrogenase complex and oxidative phosphorylation by the FbcBCF and the high-affine terminal oxidases FixNOPQ and CtaCDBBGE.

**Catabolism of malate under operational TCA**

$$\mathrm{Malate} + \mathrm{Q} + 5\,\mathrm{NAD} + \mathrm{ADP} + \mathrm{Pi} + 2\,\mathrm{H_2O} \longrightarrow \mathrm{QH_2} + 4\,\mathrm{CO_2} + 5\,\mathrm{NADH} + \mathrm{ATP} \qquad [1]$$

multiply 1 by $\frac{8}{5}$ [2]

$$1.6\,\mathrm{Malate} + 1.6\,\mathrm{Q} + 8\,\mathrm{NAD} + 1.6\,\mathrm{ADP} + 1.6\,\mathrm{Pi} + 3.2\,\mathrm{H_2O} \longrightarrow 1.6\,\mathrm{QH_2} + 6.4\,\mathrm{CO_2} + 8\,\mathrm{NADH} + 1.6\,\mathrm{ATP} \qquad [3]$$

Electron bifurcation of NADH by the FixABCX complex:

$$8\,\mathrm{NADH_2} + 8\,\mathrm{Fd_{ox}} + 4\,\mathrm{Q} \longrightarrow 8\,\mathrm{NAD} + 8\,\mathrm{Fd_{red}} + 4\,\mathrm{QH_2} \qquad [4]$$

Nitrogenase reaction by NifDK (as biochemical equation not balanced for $H^+$):

$$\mathrm{N_2} + 8\,\mathrm{Fd_{red}} \longrightarrow 2\,\mathrm{NH_3(aq)} + \mathrm{H_2} + 8\,\mathrm{Fd_{ox}}, \qquad \text{requires } 8\,\mathrm{H^+},\ 16\,\mathrm{ATP} \qquad [5]$$

Oxidative phosphorylation by FbcBCF and high-affine terminal oxidase FixNOPQ:

$$1.6\,\mathrm{QH_2} + 4\,\mathrm{QH_2} = 5.6\,\mathrm{QH_2}, \text{Sum of } \mathrm{QH_2} \text{ production by TCA 3 and electron bifurcation 4} \qquad [6]$$

$$5.6\,\mathrm{QH_2} + 2.8\,\mathrm{O_2} \longrightarrow 5.6\,\mathrm{Q} + 5.6\,\mathrm{H_2O} \qquad (5.6 \cdot 6\,\mathrm{H^+} = 33.6\,\mathrm{H^+} \text{ exported}) \qquad [7]$$

Overall biochemical reaction, not yet balanced for charge and $H^+$:

$$1.6\,\mathrm{malate} + \mathrm{N_2} + 2.8\,\mathrm{O_2} \longrightarrow 2\,\mathrm{NH_3} + 6.4\,\mathrm{CO_2} + \mathrm{H_2} + 0.8\,\mathrm{H_2O} \qquad \text{biochem. eq.} \qquad [8]$$

Balancing the reaction stoichiometry for charge and $H^+$:

$$1.6\,\mathrm{C_4H_4O_5}^{2-} + \mathrm{N_2} + 2.8\,\mathrm{O_2} \longrightarrow 2\,\mathrm{NH_4}^{+} + 6.4\,\mathrm{CO_2} + \mathrm{H_2} + 0.8\,\mathrm{H_2O} \qquad \text{chemical eq.} \qquad [9]$$

$$6.4\,\mathrm{C} = 6.4\,\mathrm{C} \qquad \text{C balanced} \qquad [10]$$

$$2\,\mathrm{N} = 2\,\mathrm{N} \qquad \text{N balanced} \qquad [11]$$

$$13.6\,\mathrm{O} = 13.6\,\mathrm{O} \qquad \text{O balanced} \qquad [12]$$

$$6.4\,\mathrm{H} \neq 11.6\,\mathrm{H} \qquad \text{5.2 Hmissing} \qquad [13]$$

$$-3.2 \neq +2 \qquad \text{+ 5.2 missing} \qquad [14]$$

Insertion of 5.2 $H^+$ to the left of 9 results in the following charge and $H^+$ corrected equation:

$$1.6\,\mathrm{C_4H_4O_5}^{2-} + \mathrm{N_2} + 2.8\,\mathrm{O_2} + 5.2\,\mathrm{H^+} \longrightarrow 2\,\mathrm{NH_4}^{+} + 6.4\,\mathrm{CO_2} + \mathrm{H_2} + 0.8\,\mathrm{H_2O} \text{chemical eq.} \qquad [15]$$

Proton translocation:

$$33.6\,\mathrm{H^+}_{\mathrm{in}} \longrightarrow 33.6\,\mathrm{H^+}_{\mathrm{out}} \qquad \mathrm{O_2} \text{ dependent proton translocation from 7} \qquad [16]$$

$$33.6\,\mathrm{H^+}_{\mathrm{out}} \longrightarrow 33.6\,\mathrm{H^+}_{\mathrm{in}} \qquad \text{protons available from 7} \qquad [17]$$

$$5.2\,\mathrm{H^+}_{\mathrm{out}} \longrightarrow 5.2\,\mathrm{H^+}_{\mathrm{in}} \qquad \text{influx of } \mathrm{H^+} \text{ derived from 15} \qquad [18]$$

$$38.8\,\mathrm{H^+}_{\mathrm{out}} \longrightarrow 38.8\,\mathrm{H^+}_{\mathrm{in}} \qquad \text{17 + 18 } \textbf{Sum of proton influx} \qquad [19]$$

Stochiometry of the proton motive force (PMF) dependent ATP synthesis:

$$\frac{[\mathrm{H^+}]}{[\mathrm{ATP}]} = \frac{10}{3} \qquad , \frac{38.8}{3.33} = 11.64 \text{ , ATP synthesized} \qquad [20]$$

Stoichiometry of Enzyme-coupled ATP synthesis:

1.6 ATP/ malate generated by succinyl-CoA synthase, 1 [21]

Sum of oxygen-dependent, oxygen-independent and Enzyme-coupled ATP synthesis - :

10.08 ATP Oxygen-dependent [22]

1.56 ATP Oxygen-independent [23]

1.6 ATP Enzyme-coupled [24]

13.42 ATP Sum [25]

-16 ATP Nitrogenase reaction [26]

**-2.76 ATP Net loss** [27]

[28]

Overall reaction of $N_2$ fixation by bacteroid:

$$1.6\,\mathrm{Malate} + \mathrm{N_2} + 2.8\,\mathrm{O_2} + 5.2\,\mathrm{H^+} + 2.76\,\mathrm{ATP} + 1.96\,\mathrm{H_2O} \longrightarrow 2\,\mathrm{NH_4}^+ + 6.4\,\mathrm{CO_2} + \mathrm{H_2}\,2.36\,\mathrm{ADP} + 2.36\,\mathrm{Pi} \quad [29]$$

With malate as the sole energy source, a net loss of -2.76 ATP occurs, therefore malate is likely not acting as the sole substrate of symbiotic nitrogen fixation.

**Model B: Succinate and operational TCA**

Conversion of succinate by TCA cycle:

$$\mathrm{Succinate} + 2\,\mathrm{Q} + 5\,\mathrm{NAD} + \mathrm{ADP} + \mathrm{Pi} + 3\,\mathrm{H_2O} \longrightarrow 2\,\mathrm{QH_2} + 4\,\mathrm{CO_2} + 5\,\mathrm{NADH} + \mathrm{ATP} \quad [30]$$

$$\text{multiply 30 by } \frac{8}{5} \quad [31]$$

$$1.6\,\mathrm{Succinate} + 3.2\,\mathrm{Q} + 8\,\mathrm{NAD} + 1.6\,\mathrm{ADP} + 1.6\,\mathrm{Pi} + 4.8\,\mathrm{H_2O} \longrightarrow 3.2\,\mathrm{QH_2} + 6.4\,\mathrm{CO_2} + 8\,\mathrm{NADH} + 1.6\,\mathrm{ATP} \quad [32]$$

Electron bifurcation of NADH by the FixABCX complex:

$$8\,\mathrm{NADH_2} + 8\,\mathrm{Fd_{ox}} + 4\,\mathrm{Q} \longrightarrow 8\,\mathrm{NAD} + 8\,\mathrm{Fd_{red}} + 4\,\mathrm{QH_2} \quad [33]$$

Nitrogenase reaction by NifDK (as biochemical equation not balanced for $H^+$):

$$\mathrm{N_2} + 8\,\mathrm{Fd_{red}} \longrightarrow 2\,\mathrm{NH_3(aq)} + \mathrm{H_2} + 8\,\mathrm{Fd_{ox}}, \qquad \text{requires } 8\,\mathrm{H^+},\ 16\ \mathrm{ATP} \quad [34]$$

Oxidative phosphorylation by FbcBCF and high-affine terminal oxidase FixNOPQ:

$$3.2\,\mathrm{QH_2} + 4\,\mathrm{QH_2} = 7.2\,\mathrm{QH_2}, \text{Sum of } \mathrm{QH_2} \text{ production by TCA 32 and electron bifurcation 33} \quad [35]$$

$$7.2\,\mathrm{QH_2} + 3.6\,\mathrm{O_2} \longrightarrow 7.2\,\mathrm{Q} + 7.2\,\mathrm{H_2O} (7.2 \cdot 6\,\mathrm{H^+} = 43.2\,\mathrm{H^+} \text{ exported}) \quad [36]$$

Overall biochemical reaction, not yet balanced for charge and $H^+$:

$$1.6\,\mathrm{succinate} + \mathrm{N_2} + 3.6\,\mathrm{O_2} \longrightarrow 2\,\mathrm{NH_3} + 6.4\,\mathrm{CO_2} + \mathrm{H_2} + 0.8\,\mathrm{H_2O} \qquad \text{biochem. eq.} \quad [37]$$

Balancing the reaction stoichiometry for charge and $H^+$:

$$1.6\,\mathrm{C_4H_4O_4}^{2-} + \mathrm{N_2} + 3.6\,\mathrm{O_2} \longrightarrow 2\,\mathrm{NH_4}^{+} + 6.4\,\mathrm{CO_2} + \mathrm{H_2} + 0.8\,\mathrm{H_2O} \qquad \text{chemical eq.} \quad [38]$$

$$6.4\,\mathrm{C} = 6.4\,\mathrm{C} \qquad \text{C balanced} \quad [39]$$

$$2\,\mathrm{N} = 2\,\mathrm{N} \qquad \text{N balanced} \quad [40]$$

$$13.6\,\mathrm{O} = 13.6\,\mathrm{O} \qquad \text{O balanced} \quad [41]$$

$$6.4\,\mathrm{H} \neq 11.6\,\mathrm{H} \qquad 5.2\,\mathrm{H^+} \text{ missing} \quad [42]$$

$$-3.2 \neq +2 \qquad +\,5.2 \text{ missing} \quad [43]$$

Insertion of 5.2 $H^+$ to the left of 38 results in the following charge and $H^+$ corrected equation:

$$1.6\,\mathrm{C_4H_4O_4}{-}2 + \mathrm{N_2} + 2.8\,\mathrm{O_2} + 5.2\,\mathrm{H^+} \longrightarrow 2\,\mathrm{NH_4}^{+} + 6.4\,\mathrm{CO_2} + \mathrm{H_2} + 0.8\,\mathrm{H_2O} \qquad \text{chemical eq.} \quad [44]$$

Proton translocation:

$$43.2\,\mathrm{H^+}_{\mathrm{in}} \longrightarrow 43.2\,\mathrm{H^+}_{\mathrm{out}} \qquad \mathrm{O_2} \text{ dependent proton translocation from 7} \quad [45]$$

$$43.2\,\mathrm{H^+}_{\mathrm{out}} \longrightarrow 43.2\,\mathrm{H^+}_{\mathrm{in}} \qquad \text{protons available from 7} \quad [46]$$

$$5.2\,\mathrm{H^+}_{\mathrm{out}} \longrightarrow 5.2\,\mathrm{H^+}_{\mathrm{in}} \qquad \text{influx of } \mathrm{H^+} \text{ derived from 44} \quad [47]$$

$$48.8\,\mathrm{H^+}_{\mathrm{out}} \longrightarrow 48.8\,\mathrm{H^+}_{\mathrm{in}} \qquad 46 + 47\ \textbf{Sum of proton influx} \quad [48]$$

Stochiometry of the proton motive force (PMF) dependent ATP synthesis:

$$\frac{[\mathrm{H^+}]}{[\mathrm{ATP}]} = \frac{10}{3}, \quad \frac{48.8}{3.33} = 14.64, \text{ ATP synthesized} \quad [49]$$

Stoichiometry of Enzyme-coupled ATP synthesis:

1.6 ATP generated by Enzyme-coupled ATP synthesis of TCA [50]

Sum of oxygen-dependent, oxygen-independent and Enzyme-coupled ATP synthesis - :

12.96 ATP Oxygen-dependent [51]

1.56 ATP Oxygen-independent [52]

1.6 ATP Enzyme-coupled [53]

16.12 ATP Sum [54]

-16 ATP Nitrogenase reaction [55]

**0.12 ATP Net gain** [56]

Overall reaction of $N_2$ fixation by bacteroid:

$$1.6\,\mathrm{succinate} + \mathrm{N_2} + 3.6\,\mathrm{O_2} + 5.2\,\mathrm{H^+} + 0.12\,\mathrm{ADP} + 0.12\,\mathrm{Pi} \longrightarrow 2\,\mathrm{NH_4}^{+} + 6.4\,\mathrm{CO_2} + \mathrm{H_2} + 0.12\,\mathrm{ATP} + 0.92\,\mathrm{H_2O} \quad [57]$$

With Succinate as the sole energy source, net production of 0.12 ATP occurs. However, net ATP gain does not occur if more than 0.4 protons out of a total of 48.8 per $N_2$ fixed are transported by membrane diffusion rather than by the action of ATP synthase. Over 99.1% of the protons generated must pass the ATP synthase complex, otherwise, no ATP gain is achieved. Such efficiency is likely not realistic for a biochemical process giving the high permeability of the biological cell membrane for protons.
